# Fc receptor-like 6 (FCRL6) defines pre-BCR dependent and independent pathways of natural antibody selection

**DOI:** 10.1101/667345

**Authors:** Kazuhito Honjo, Woong-Jai Won, Rodney G. King, Lara Ianov, David K. Crossman, Juliet L. Easlick, Mikhail A. Shakhmatov, Mohamed Khass, Andre M. Vale, Robert P. Stephan, Ran Li, Randall S. Davis

## Abstract

B-1a cells produce ‘natural’ antibodies (Abs) to neutralize pathogens and clear neo self-antigens, but the fundamental selection mechanisms that shape their polyreactive repertoires are poorly understood. Here, we identified a B cell progenitor subset defined by Fc receptor-like 6 (FCRL6) expression, harboring innate-like defense, migration, and differentiation properties conducive for natural Ab generation. Compared to FCRL6^-^ pro B cells, the repressed mitotic, DNA damage repair, and signaling activity of FCRL6^+^ progenitors, yielded V_H_ repertoires with biased distal *Ighv* segment accessibility, constrained diversity, and hydrophobic and charged CDR-H3 sequences. Beyond nascent autoreactivity, V_H_11 productivity, which predominates phosphatidylcholine-specific B-1a B cell receptors (BCRs), was higher for FCRL6^+^ cells as was pre-BCR formation, which was required for Myc induction and V_H_11, but not V_H_12, B-1a development. Thus, FCRL6 revealed unexpected heterogeneity in the developmental origins, regulation, and selection of natural Abs at the pre-BCR checkpoint with implications for autoimmunity and lymphoproliferative disorders.

## Introduction

B-1 B cells, and specifically B-1a cells that express CD5, are the primary source of ‘natural’ antibodies (Abs) whose poly/autoreactive features provide homeostatic protection against bacterial and viral pathogens (Ochsenbein et al., 1999; Baumgarth et al., 2005; Baumgarth, 2011). Natural Abs also serve housekeeping functions by clearing apoptotic cells and neo self-antigens (Ags) (Shaw et al., 2000). B-1a cells home to the spleen, as well as the peritoneal (PeC) and pleural cavities (Baumgarth, 2011; Montecino-Rodriguez and Dorshkind, 2012), where a remarkably large proportion (∼5-15%) express stereotypic B cell receptors (BCRs) restricted to V_H_11 and V_H_12, which both recognize phosphatidylcholine (PtC), a determinant present in cell membranes and certain bacteria (Mercolino et al., 1988; Sohlenkamp et al., 2003; Rowley et al., 2007). Consequently, the Ab repertoires of B-1a cells have a biased composition. Due to a lack of junctional diversity during fetal life (Feeney, 1990), they generate germline-related Abs encoding CDR-H3 segments that are more hydrophobic than B-2 repertoires (Khass et al., 2018). Splenic marginal zone (MZ) B cells similarly possess innate-like defense properties and fetal-related V_H_ repertoires enriched in charged CDR-H3 segments (Martin and Kearney, 2002; Carey et al., 2008; Khass et al., 2018). The propensity to generate autoreactive Abs has thus implicated innate-like MZ and B-1 B cells in the pathogenesis of autoimmunity (AI) and malignancy, including the most common leukemia in Western countries, chronic lymphocytic leukemia (CLL) (Gu et al., 1990; Chiorazzi and Ferrarini, 2011; Hayakawa et al., 2016).

The extrafollicular localization and constrained repertoires of B-1 cells are in marked contrast to B-2 cells that participate in T cell-dependent responses in secondary lymphoid tissues and generate affinity-matured, highly-diversified Abs. However, the developmental origins and regulatory mechanisms governing the selection of B-1 cells remain debated. The “lineage” hypothesis, proposes that B-1 cells differentiate from discrete progenitors chiefly during fetal development and early ontogeny (Herzenberg and Herzenberg, 1989; Montecino-Rodriguez et al., 2006; Montecino-Rodriguez and Dorshkind, 2012). Alternatively, the “selection” hypothesis posits that the Ag reactivity of the BCR mediates lineage specification and compartmentalization (Haughton et al., 1993; Arnold et al., 2000).

B-1a B cells mainly derive from the fetal liver (FL) and neonatal tissues, but can develop, albeit less efficiently, from adult bone marrow (BM) where B-2 differentiation predominates (Montecino-Rodriguez et al., 2006; Baumgarth, 2011; Montecino-Rodriguez and Dorshkind, 2012). A temporal and anatomic switch in primary B cell development is mediated by the Let7-Lin28b-Arid3a axis that differentially regulates B-1 development during fetal versus adult life (Yuan et al., 2012; Zhou et al., 2015; Kristiansen et al., 2016). Although several transcription factors can preferentially influence the B-1 pathway, *Bhlhe41* has a distinct impact on B-1a differentiation, self-renewal, and BCR repertoire formation (Kreslavsky et al., 2017). However, a model linking the cellular origins with selection mechanisms that generate and shape the characteristic poly/autoreactive repertoires of B-1a cells is still lacking.

Based on conventional B-2 selection models, the pre-BCR establishes a critical checkpoint to promote the emergence of IgM heavy chains (μHC) with tyrosine-enriched CDR-H3 rearrangements, rather than charged or hydrophobic loops that could harbor self-reactivity (Bankovich et al., 2007; Keenan et al., 2008; Khass et al., 2016). Hence, Hardy postulated that μHCs from natural Abs, and V_H_11 in particular (Wasserman et al., 1998; Rowley et al., 2007), pair less efficiently with the surrogate light chain (SLC) concluding that pre-BCR selection likely differs during the fetal versus adult stages of ontogeny. Indeed, V_H_11 interactions with the SLC appear weaker than other V_H_ segments (Yoshikawa et al., 2009). Alternatively, B-1 progenitor μHCs, including V_H_12 segments, may instead bypass pre-BCR selection by pairing with prematurely rearranged light chains (Wong et al., 2017).

Here we found that expression of Fc receptor-like 6 (FCRL6) defined subpopulations of B cell progenitors throughout ontogeny that reflect fetal versus adult B-1a developmental potential. FCRL6^+^ FL and BM pro B cells exhibited protracted differentiation and proliferation, including the generation of nascent μHCs harboring constrained diversity and autoreactive properties. Furthermore, FCRL6 discriminated pre-BCR dependent and independent selection pathways that differentially impacted V_H_11 and V_H_12 B-1a cell development. FCRL6^+^ progenitors exhibited attributes of TCF/LEF and nervous system developmental regulation as well as B-1a related defense, migration, and differentiation properties. These findings provide novel insight into the heterogeneous origins and selection mechanisms underlying B-1a development and have implications for AI and CLL pathogenesis.

## Results

### FCRL6 segregates subsets of B cell progenitors throughout ontogeny

Following the discovery of human *FCRL1-6* (Davis et al., 2001; Davis, 2007), we identified a mouse *Fcrl6* counterpart encoding putative type I transmembrane (TM), glycosylphosphatidylinositol (GPI)-linked, and secreted isoforms with two extracellular Ig-like domains (Davis, 2007). By RQ-PCR, we detected *Fcrl6* transcripts in primary lymphopoietic tissues, including embryonic day 18 (E18) FL, day 3 postnatal liver, and adult BM (Figure 1A). The generation of monoclonal Abs (mAbs) yielded two FCRL6-specific subclones (1C3 and 3C1) and identified that the TM isoform had a molecular weight of ∼42kD (Figure supplement 1A,B). Although FCRL6 is confined to natural killer (NK) and T cells in humans (Schreeder et al., 2008), by flow cytometry analysis from adult BALB/cJ BM, FCRL6 was restricted to early stage B cells (Figure supplement 1C and data not shown). Expression was not detected by other lineages or in any other hematopoietic tissue at homeostasis. FCRL6 shared a similar pattern of expression by FL and BM B cells bearing CD19 and B220 as well as AA4.1 and CD43, but little if any surface IgM (Figure supplement 2A). These observations indicated that 11-12% of FL and 1-2% of BM CD19^+^ B cells express FCRL6.

**Figure 1.**
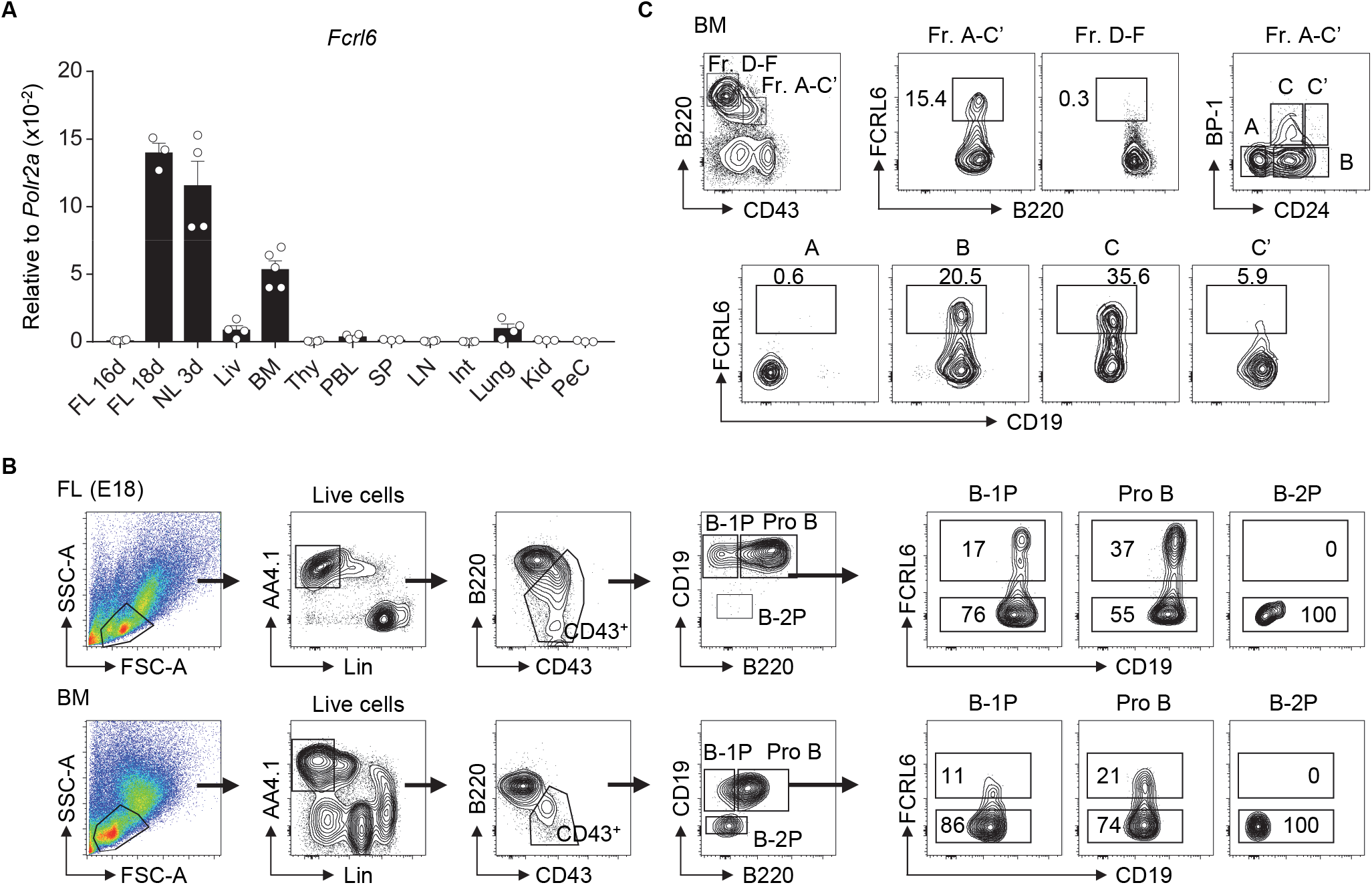
*Fcrl6* is restricted to primary B lymphopoietic tissues and segregates B lineage progenitors. (**A**) RQ-PCR analysis of *Fcrl6* expression in BALB/cJ tissues including embryonic day 16 (E16) and 18 (E18) fetal liver (FL), 3 day neonatal liver (NL), and adult (all others) liver (Liv), bone marrow (BM), thymus (Thy), peripheral blood leukocytes (PBL), spleen (SP), lymph node (LN), small intestine (Int), lung, kidney (Kid), and peritoneal cavity lavage cells (PeC). Primers were specific for regions of exons 1-3 encoding the signal peptide and first extracellular domain. Samples were normalized to *RNA polymerase II* (*Polr2a*) expression. (**B**) Flow cytometry with a receptor-specific mAb (1C3) analyzing FCRL6 surface expression by B-1 progenitor (B-1P), B-2 progenitor (B-2P) and pro B cell subpopulations in E18 FL and adult BM according to the scheme of Montecino-Rodriguez et al., 2006. Numbers within the gates indicate the percentage of cells. (**C**) Flow cytometry analysis of adult BM B cell fractions (Fr) according to the differentiation scheme of Hardy et al., 2001. Numbers adjacent to the gates indicate the percentage of cells. Data shown are cumulative from the tissues of 3-5 mice amplified in duplicate (**A**), representative of more than five independent experiments (**B**), or three independent experiments (**C**).

Because its frequency was about 10 times greater in the FL, we examined FCRL6 using a differentiation scheme that segregates precursors capable of reconstituting the B-1 or B-2 lineage (Montecino-Rodriguez et al., 2006). FCRL6 marked subsets of CD19^hi^B220^hi^ pro B cells in the FL and BM, as well as some CD19^hi^B220^lo^ B-1 precursors (B-1P), which are more frequent in the FL, but not CD19^lo^B220^+^ B-2 precursors (B-2P) that chiefly populate the BM (Figure 1B). In day 3 and 7 neonatal tissues, FCRL6^+^ B cells variably expanded in frequency with time in the liver, BM, and spleen (Figure supplement 2B). This pattern of tissue expression demonstrated that FCRL6 segregates subsets of progenitor B cells conserved throughout life.

We then examined adult BM according to Hardy’s definition (Hardy and Hayakawa, 2001). By this scheme, only a minor subset of B cell progenitors, estimated at 7-15% of the total A-C’ fraction (Fr), expressed FCRL6 (Figure 1C). Further separation based on CD24/HSA and BP-1 disclosed hardly any FCRL6^+^ Fr. A cells, but the frequency increased in Fr. B, peaked in Fr. C, declined in Fr. C’, and very few cells were found beyond the Fr. D stage. Thus, FCRL6 expression was tightly regulated and predominantly restricted to Fr. B and C pro and pre B cells, developmental stages during which V(D)J recombination and pre-BCR selection occur.

### FCRL6 progenitors exhibit features of protracted differentiation

We next analyzed other phenotypic features in the FL and BM. By forward scatter, FCRL6^+^ pro B cells were smaller than FCRL6^-^ cells, suggesting they might proliferate more slowly (Figure 2A). The relatively lower density of CD127/IL-7Rα on FCRL6^+^ cells in both tissues indicated they might also be less responsive to IL-7. The early differentiation marker CD117/c-kit was uniformly upregulated by the FCRL6^+^ subset in FL, but not in the BM. BP-1 was also less abundant on FL FCRL6^+^ cells. Finally, MHCII and CD138 were slightly higher on FCRL6^-^ pro B cells in the BM than FL. Together these results indicated that FCRL6 distinguished less differentiated pro B cells.

**Figure 2.**
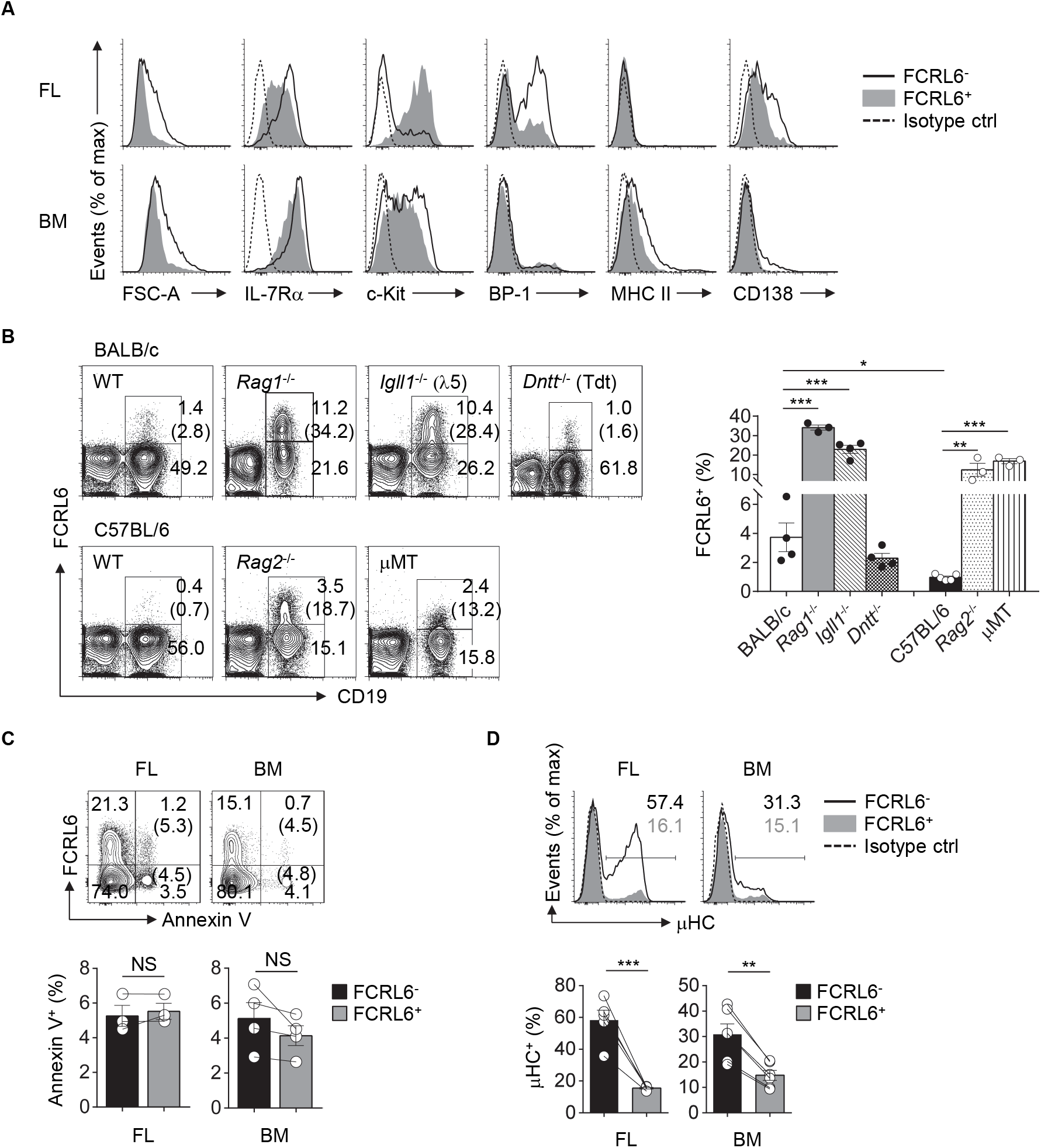
FCRL6^+^ pro B cells exhibit protracted development, expand in *Igh* defective mice, and generate little cytoplasmic IgM (μHC). (**A**) Flow cytometry analyzing FCRL6^+^ and FCRL6^-^ pro B cells in E18 FL (above) and adult BM (below) to assess cell size by forward scatter (FSC) and co-expression of indicated surface markers. (**B**) Flow cytometry of total BM cells isolated from adult WT BALB/c and C57BL/6 mice or genetic mutants (*n* = 3-5 per strain) on related background strains stained for FCRL6 and CD19 as well as IgM to dump immature/mature B cells. Numbers adjacent to the gates indicate the percent of cells among total lymphocytes. The frequency of FCRL6^+^/CD19^+^ B cells is indicated in parentheses and summarized in the bar graph to the right. (**C**) Survival differences of FL and BM FCRL6^+^ and FCRL6^-^ pro B cells evaluated by Annexin V staining and flow cytometry. The frequency of FCRL6^+^ or FCRL6^-^ Annexin V^+^ pro B cells is indicated in parentheses. (**D**) Representative intracellular IgM staining of FL and BM FCRL6^-^ and FCRL6^+^ pro B cells. Each symbol represents an individual mouse. Small horizontal lines (**B**-**D**) indicate the mean (± s.e.m.). NS, not significant (*P* > 0.05); \**P* < 0.05; \*\**P* < 0.01; \*\*\**P* < 0.001 as determined by unpaired (**B**) or paired (**C, D**) Student’s *t*-test. Data are representative of at least two experiments (**A**-**C**), or more than 10 independent experiments (**D**).

We then compared different mouse strains and models with defects in early B cell development. We first observed that the frequency of CD19^+^FCRL6^+^ cells in the BM was about four-fold higher in WT BALB/c than C57BL/6 mice (Figure 2B). In *Rag1*^-/-^*, Rag2*^-/-^, and *Igll1* (λ5)*^-/-^* mutants, B cell development is blocked at Fr. C and C’ resulting in the enrichment of Fr. B and C pro B cells. BM cells from *Rag1*^-/-^*, Rag2*^-/-^, *Igll1^-/-^*, and μMT mice of respective strains all had expanded frequencies of FCRL6^+^ B cells. In these models, CD19^+^FCRL6^+^ B cells were ∼10-15 fold greater compared to WT mice. However, no difference was evident in *Tdt*^-/-^ mice. Thus, FCRL6^+^ cells emerged prior to RAG1/2 expression, and moreover V(D)J recombination, segregated a subset of pro B cells that varied by genetic background, and expanded following disruption of *Igh* rearrangement or pre-BCR/BCR assembly.

Because FCRL6^+^ pro B cells were relatively smaller and less differentiated, it was possible they represented defective cells destined for elimination. However, no differences in Annexin V reactivity, and thus survival status, were found (Figure 2C). We then analyzed their capacities for generating cytoplasmic μHC. Compared to FCRL6^-^ progenitors, frequencies of μHC^+^FCRL6^+^ cells were nearly four-fold lower in the FL and two-fold lower in the BM (Figure 2D). These results indicated marked mechanistic differences in μHC generation according to FCRL6 status.

### FCRL6 defines distinct transcriptomic and biologic heterogeneity

We next performed high-throughput RNA sequencing (RNA-seq) to compare the gene expression profiles of FL and BM FCRL6^+^ to FCRL6^-^ pro B cells. By principle component analysis (PCA), duplicate samples clustered closely and segregated into four quadrants indicating distinct gene signatures for the four subsets (Figure 3A). This analysis identified a total of 1,276 differentially expressed genes (DEGs) in the FL and 384 in the BM (DEG criteria: a change in expression of one-fold (log_2_ value); false discovery rate (FDR), <0.05) (Figure 3B - Figure supplement 3A,B). By gene ontogeny (GO) analysis, cytokine production, adhesion, innate defense, and migration pathways were upregulated by FL FCRL6^+^ pro B cells (Figure 3C). Because these features were evocative of innate-like B cells, we compared DEGs from FL pro B cells with those from PeC B-1a and B-1b cells or splenic B-1a and MZ B cells extracted from the ImmGen database. Greater gene overlap (∼1.5 fold) was observed with FCRL6^+^ upregulated DEGs (Figure supplement 3C). Distinct biologic differences related to FCRL6 expression were further implied by 265 DEGs that overlapped between the FL and BM (Figure 3B,D). Dissimilar proliferation kinetics in FCRL6^+^ cells were implied by the strong repression of mitotic cell cycle genes, including transcripts for *Mki67* (encodes Ki-67). Intracellular staining confirmed higher frequencies of Ki-67^+^ FCRL6^-^ pro B cells (Figure 3E). Furthermore, an analysis of BrdU and 7-AAD status showed that most FCRL6^+^ cells had not entered the S/G2+M phase and were primarily in G0/G1 (Figure 3F). G2/M frequencies were 6.5 and 5.7 fold higher for FCRL6^-^ compared to FCRL6^+^ pro B cells in the FL and BM. These results demonstrated significantly lower proliferation and cell cycle activity for FCRL6^+^ cells *in vivo* regardless of their tissue of origin. By GO pathway analysis, genes for signal transduction regulation were also induced. Among DEGs encoding phosphatases, 14/20 in the FL and 4/5 in the BM were upregulated in FCRL6^+^ cells (Figure supplement 3D). We thus compared signaling features of sorted BM pro B cells by *ex vivo* treatment with the phosphatase inhibitor pervandate. FCRL6^-^ cells had higher STAT5 and ERK phosphorylation (Figure 3G), indicating the diminished activation potential of FCRL6^+^ cells. Thus, in addition to other properties, FCRL6 defined pro B subsets with specific regulatory differences.

**Figure 3.**
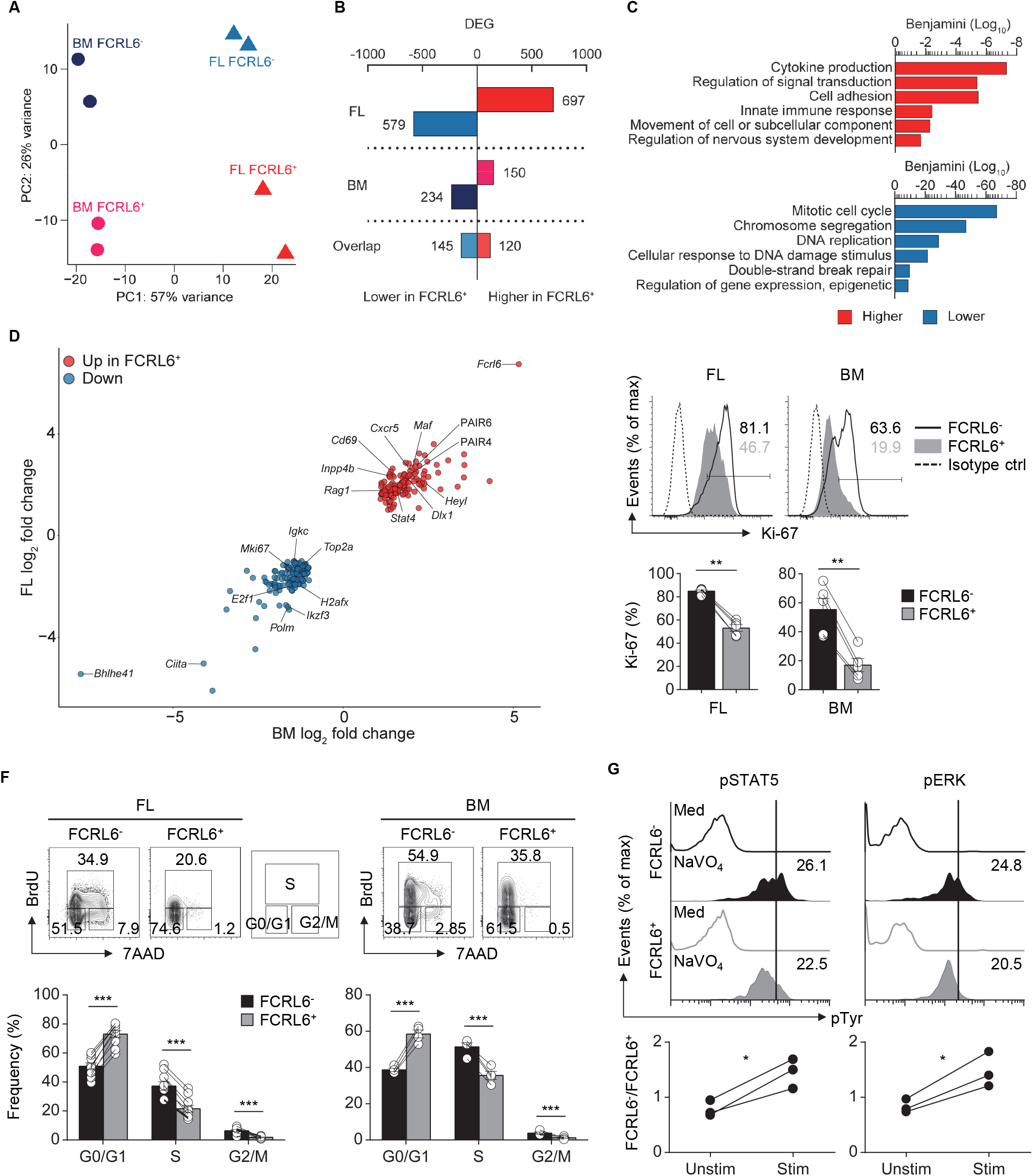
Transcriptomic and biologic heterogeneity of pro B cells. (**A**) Principal component (PC) analysis of RNA-seq performed on duplicate FL and BM FCRL6^+^ and FCRL6^-^ pro B cell samples sorted by flow cytometry as in Figure 1B. (**B**) Quantitation of differentially expressed genes (DEG) relative to FCRL6^+^ cells by tissue defined by a change in expression of one-fold (log_2_ value) and FDR (<0.05). (**C**) Gene ontogeny (GO) pathways derived from FL DEGs. (**D**) Scatter plot of overlapping DEGs. Results are presented as the difference in expression (log_2_ transformed values) of FCRL6^+^ relative to FCRL6^-^ pro B cells from the FL (vertical) and BM (horizontal). (**E**) Intracellular staining and flow cytometry analysis of Ki-67 among pro B cell subsets (*n* = 5 each). Percentages of gated cells relative to an isotype-matched control are indicated in the histograms and summarized below. (**F**) Proliferation and cell cycle status of FL (*n* = 10) and BM (*n* = 4) pro B cell subsets assayed by staining with the thymidine analog BrdU and 7-AAD DNA intercalation dye before flow cytometry analysis. The percentages of G0/G1, S, and G2/M phase cells are shown adjacent to the contour plot gates and are summarized below. (**G**) Phospho (p)-flow cytometry analysis of sorted FCRL6^+^ and FCRL6^-^ BM pro B cells either unstimulated (medium) or stimulated with sodium pervanadate (NaVO_4_) for 10 min *ex vivo*. Cells were intracellularly stained with control Abs or those specific for pSTAT5 and pERK. Values in the histograms represent MFI ratios (test Ab MFI/control Ab MFI). Fold differences comparing FCRL6^-^ and FCRL6^+^ pro B cells are shown below. Each symbol (**E-G**) represents an individual mouse. Small horizontal lines (**E, F**) indicate the mean (± s.e.m.). \**P* < 0.05; \*\**P* < 0.01; \*\*\**P* < 0.001 as determined by paired Student’s *t*-test. Data are from four independent sorts and two per cell type (**A**-**D**), one representative of two experiments (**E**-**F**), and three independent experiments (**G**).

As expected, *Fcrl6* was the most upregulated overlapping DEG between tissues. Remarkably, *Bhlhe41*, a transcriptional repressor critical for B-1a development and Ab repertoire formation (Kreslavsky et al., 2017), proved the most downregulated (Figure 3D). Such polarization suggested that FCRL6 was a defining developmental marker. Furthermore, the upregulation of GO pathway components associated with nervous system development, including overlapping DEGs (*Heyl*, *Dlx1*, *Aph1b*) and pathways relevant for Notch, Wnt, stem cell, EMT biology, and TCF/LEF (Figure supplement 3E), collectively demonstrated divergent developmental and regulatory features for FCRL6^+^ progenitors.

### Disparate DNA repair, *Ighv* accessibility and μHC repertoires

Because V(D)J recombination and DNA damage repair are coupled to the cell cycle (Lin and Desiderio, 1995; Bassing and Alt, 2004), enrichment in the G0/G1 phase indicated that FCRL6^+^ pro B cells might be more actively undergoing HC rearrangement. Indeed, the RNA-seq GO pathway analysis revealed the downregulation of DNA damage repair programs (Figure 3C). To assess the spectrum of transcription, we grouped the *Ighv* locus into four domains according to the locations of CTCF binding sites that organize locus contraction and transcript accessibility during V(D)J recombination (Choi et al., 2013) (Figure 4A). PCA of normalized *Ighv* segments showed partitioning of duplicate samples into quadrants, but the variance between BM sets was smaller than that of FL samples (Figure supplement 4A). By unsupervised Euclidian clustering, we found that the *Ighv* signature of FL FCRL6*^+^* pro B cells segregated independently, suggesting that this subset generated a distinct complement of V_H_ segments (Figure supplement 4B). FCRL6^+^ cells primarily upregulated distal *Ighv1*/J558 segments from domain 4, but downregulated proximal domain 1 and 2 *Ighv* genes (Figure 4A-C). Indeed, multiple V_H_ segments from these domains were among FL DEGs (Figure 4D). A similar trend of differential locus accessibility was evident in the BM, but did not reach the same threshold of significance (data not shown). Notably, no differences were evident for *Ighv5-2*/V_H_81X or segments encoding PtC-reactive natural Abs, *Ighv12-3* and *Ighv11-2*. Of two phosphorylcholine (PC)-reactive HCs, *Ighv7-1* was downregulated in FCRL6^+^ cells, but *Ighv7-3* was not.

**Figure 4.**
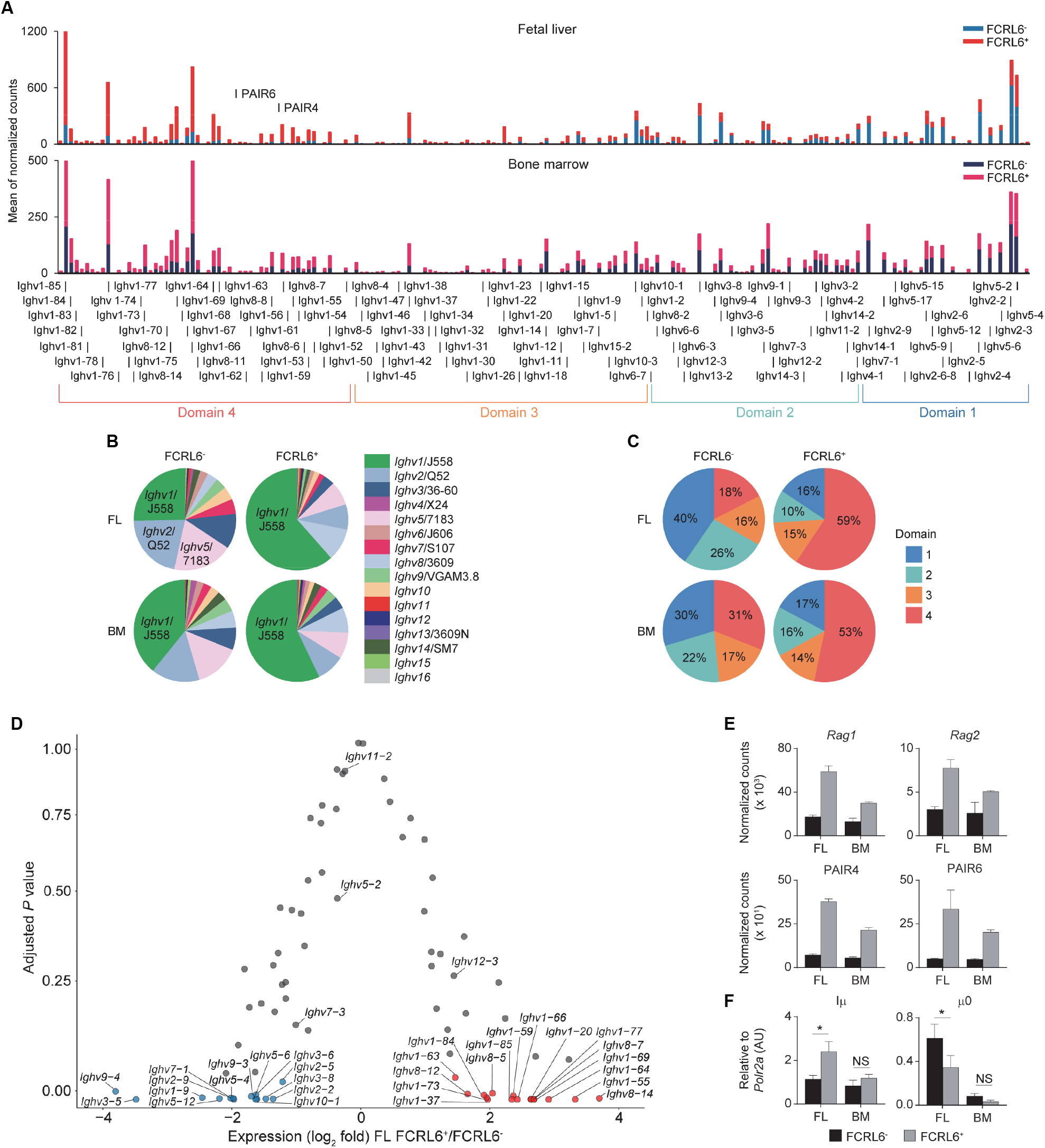
FCRL6 discriminates pro B cells with divergent *Ighv* locus accessibility. (**A**) Individual segment usage across the *Ighv* locus based on the means of normalized counts from duplicate FL and adult BM pro B cell subsets. The locus is segregated into four domains based on Choi et al., 2013. The PAIR4 (ENSMUSG00000104373/ Gm37511) and PAIR6 (ENSMUSG00000104373/Gm37511-Gm30996) ncRNA elements are indicated in domain 4. (**B**) Frequency of *Ighv* family usage among the four subsets based on the summation of the means of normalized counts from total *Ighv* segments of each subfamily. (**C**) Frequency of domain usage based on the summation of the means of normalized counts from grouped *Ighv* locus genes segregated as indicated in **A**. (**D**) Volcano plot detailing RNA-seq analysis of *Ighv* genes from sorted FL FCRL6^+^ and FCRL6^-^ pro B cells. Results are presented as the difference in fold expression (log_2_ transformed values) of FCRL6^+^ relative to FCRL6^-^ pro B cells and plotted against adjusted *P* values. (**E**) RNA-seq analysis of coding and noncoding genes relevant to V(D)J rearrangement and locus contraction from the four FL and BM FCRL6^+^ and FCRL6^-^ pro B cell subsets. Note *Rag2* is among FL DEGs, while the other DEGs overlap between the FL and BM. (**F**) RQ-PCR of noncoding transcripts involved in initiating locus recombination relative to *Polr2a*. Small horizontal lines (**E-F**) indicate the mean (± s.e.m.) \**P* < 0.05 as determined by paired Student’s *t*-test. Data are from four independent sorts and two per cell type as in Figure 3 (**A**-**E**) and three independent sorts per subset performed in duplicate (**F**).

We next examined how *Ighv* accessibility was differentially regulated. First, recombinatorial activity was elevated. FCRL6^+^ pro B cells demonstrated upregulation of both *Rag1* and *Rag2* transcripts by RNA-seq (Figure 4E). Second, non-coding RNAs (ncRNA) critical for *Igh* locus contraction were investigated (Stubbington and Corcoran, 2013). Among the 14 PAX5-associated intergenic repeat (PAIR) elements that serve as CTCF, PAX5, and E2A binding sites (Ebert et al., 2011), ncRNA transcripts for PAIR4 and PAIR6, which are distally positioned in domain 4, were among upregulated, overlapping DEGs in FCRL6^+^ cells (Figure 3D and Figure 4E). Within the proximal J_H_-Cμ region, Iμ transcripts initiate from the Eμ intronic enhancer (Lennon and Perry, 1985) and form long-range DNA loops that tether proximal and distal ends of the *Igh* locus via direct binding to both PAIR4 and PAIR6 in a YY1-dependent fashion (Guo et al., 2011; Verma-Gaur et al., 2012). μ0 transcripts arise just upstream of the most 3’ D_H_ gene (DQ52) (Thompson et al., 1995). RQ-PCR indicated higher Iμ, but lower μ0 transcripts in FCRL6^+^ FL pro B cells. With respect to accessibility factors, transcripts for *Spi1* (PU.1), *Tcf3* (E2A), *Ikzf1* (Ikaros)*, Ikzf3* (Aiolos)*, Ctcf*, *Rad21,* and *Ezh2* were downregulated, whereas *Ebf1* was upregulated, but *Pax5* and *Yy1* did not differ (Figure supplement 4C). Collectively, these data indicated that FCRL6 expression marked disparate regulation of *Ighv* accessibility that may relate to Eμ-dependent (long-range) versus independent (local) mechanisms of locus contraction (Guo et al., 2011; Stubbington and Corcoran, 2013).

### FCRL6^+^ cells harbor constrained repertoire diversity and CDR-H3 autoreactivity

To investigate *Ighv* sequences from these pro B cells, V(D)J rearrangements were amplified from the four sorted populations. After annotating the amplicons using the IMGT database, we identified 317,702 unique dereplicated sequences that included representation of 8/15 V_H_ families (Figure supplement 5A-C). The most striking finding was the low frequency of productive rearrangements among unique *Ighv* sequences generated from FCRL6^+^ versus FCRL6^-^ pro B cells in the FL (31% vs 79%) (Figure 5A). In the BM, where Tdt is operative (Gilfillan et al., 1993), productivity rose for cells marked by FCRL6, but was still lower than FCRL6^-^ pro B cells. Notably, *Dntt* (Tdt) expression did not differ according to FCRL6 status by RNA-seq. Among amplified V_H_ families, the frequency of total productive sequences by FCRL6^+^ cells was highest for *Ighv1*/J558 (42.6% in FL and 51.1% in BM) (Figure supplement 5C). However, the frequency of productive *Ighv1*/J558 rearrangements in FCRL6^+^ cells was very low in FL, but increased in BM (Figure supplement 5B, top). Surprisingly, the most productive FL FCRL6^+^ cell rearrangements derived from the small *Ighv11* family (49.5%). However, *Ighv11-2* productivity in the BM was similar for both subsets. *Ighv5-2/*V_H_81X was the most proficient segment among FL FCRL6^+^ cells (70.8%), but was similar in FCRL6^-^ pro B cells (70.3%). However, V_H_81X productivity globally dropped in the BM. Among D and J segments, productivity for FCRL6^+^ cells was higher for D-less joins in the FL and *Ighd1*/DFL16.1 in the BM, whereas *Ighd2*/DSP segments were generally less favored. *Ighj1-4* segment usage did not markedly differ by FCRL6 status. The CDR-H3 diversity of FCRL6^+^ cell rearrangements was relatively lower in both tissues (Figure 5B). This was coincident with generally decreased CDR-H3 length and increased J trimming in the FL (Figure supplement 5B). Hence, the generally lower productivity and diversity of V(D)J rearrangements reflected significant mechanistic differences according to FCRL6 expression.

**Figure 5.**
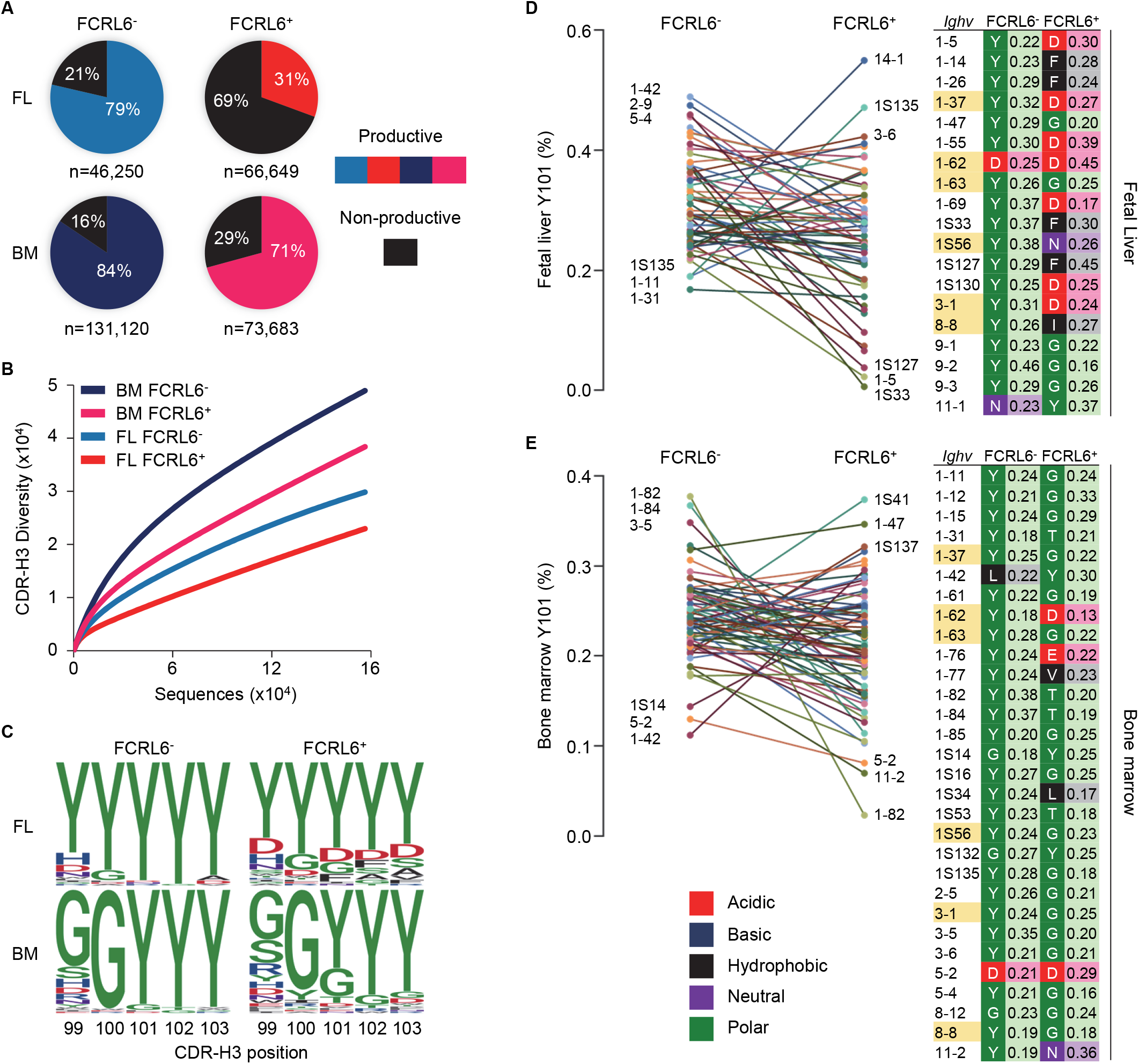
Constrained μHC productivity and repertoire diversity yields hydrophobic and charged CDR-H3s. μHC V(D)J sequences from FCRL6^+^ and FCRL6^-^ pro B cells, sorted from pooled FL E18 or adult BM BALB/c mice by flow cytometry as in Figure 1B, were amplified using 8 V_H_ family consensus and IgM-specific primers, sequenced by Illumina MiSeq, and analyzed with IMGT and R-based software. (**A**) Productivity of total dereplicated unique rearrangements from four FL and BM FCRL6^+^ and FCRL6^-^ pro B cell subsets. Frequencies are indicated within each pie chart and numbers of sequences analyzed per subset are indicated below. (**B**) Rarefaction analysis of CDR-H3 segment diversity defined by Shannon index for total replicated sequences. Numbers of sequences analyzed are listed in Figure supplement 5A. (**C**) Logo plots indicating the probability of amino acid usage at positions 99-103 of the CDR-H3 according to PDB numbering of encoded sequences (sequence numbers, Figure supplement 6A). Amino acid biochemical characteristics (color coded) are indicated in **E**. (**D**-**E**) Comparisons of tyrosine usage frequency at CDR-H3 position 101 (Y101) for 60 FL (**D**) and 74 BM (**E**) *Ighv* genes with >50 unique productive sequences common between FCRL6^+^ and FCRL6^-^ pro B cell subsets. The top and bottom three *Ighv* segments are indicated for each subset. A list of *Ighv* genes (right) that predominantly harbor non-Y101 amino acids (FL, *n* = 19 and BM, *n* = 30) are shown according to subset and tissue along with their biochemical characteristics (color-coded) and relative frequencies. *Ighv* segments (*n* = 6) highlighted yellow were common to both tissues.

At the pre-BCR checkpoint, the SLC proteins, λ5 and VpreB, make extensive contacts with the CDR-H3 of the nascent μHC by forming a ‘sensing site’ to survey its biochemical properties (Bankovich et al., 2007). Recent work has demonstrated that tyrosine enrichment at position 101 of the CDR-H3, which interacts with three residues in VpreB, favors positive selection and checkpoint passage (Khass et al., 2016). We thus compared unique productive CDR-H3 sequences from the four subsets by determining amino acid usage at positions 99-103 relative to the cysteine at position 96 (PDB numbering). While variability was evident between tissues, tyrosine enrichment at these five positions was highest for FCRL6^-^ pro B cells (Figure 5C - Figure supplement 6A). In contrast, tyrosine content at residue 101 (Y101) was about 1/3 lower for FCRL6^+^ CDR-H3 sequences.

We then inspected the CDR-H3 Y101 frequency by comparing shared *Ighv* segments between FCRL6^+^ and FCRL6^-^ pro B cells with at least 50 productive sequences each. Marked differences were identified in the range of Y101 usage. Among 60 common FL *Ighv* genes, a comparably narrower spectrum of Y101 composition was evident for FCRL6^-^ pro B cell rearrangements (16.8-48.9%) versus FCRL6^+^ cells (0.6-54.9%) (Figure 5D). This trend was also apparent for 74 *Ighv* genes shared for the BM (Figure 5E). We then analyzed common sets of *Ighv* genes to assess amino acid predominance at the CDR-H3 101 position. This breakdown identified 19/60 *Ighv* genes in the FL that preferentially utilized non-Y101 amino acids (Figure 5D, right). Surprisingly, while only two FCRL6^-^ *Ighv* sequences exhibited this feature (*Ighv1-62* and *Ighv11-1*), non-Y101 usage was evident for 18 FCRL6^+^ *Ighv* genes. In the BM, 30/74 common *Ighv* genes were predominantly non-Y101 (Figure 5E, right). Although five of these were FCRL6^-^ derived, 27 came from FCRL6^+^ cells. Additionally, six non-Y101 enriched *Ighv* genes were shared between FL and BM FCRL6^+^ cells. These findings indicated that the CDR-H3 features of FCRL6^+^ pro B cells were disproportionately hydrophobic, charged, and less tyrosine enriched. Hence, FCRL6 marked progenitors predisposed to generating rearrangements with altered categories of diversity akin to the characteristics of Abs enriched in innate-like B cells (Khass et al., 2018).

We next considered *Ighv* sequence data from B-1a cells recently analyzed by the Herzenberg group (Yang et al., 2015) and CLL clones from different mouse models and strains (Figure supplement 6A). Among 150 B-1a and 291 CLL *Ighv* sequences, *Ighv1*/J558 segments were the most common and mainly domain 4 derived (Figure supplement 6B,C). *Ighv11* was the second most favored among B-1a cells and fourth among CLL clones. CDR-H3 analyses disclosed that, like FCRL6^+^ pro B cells, tyrosine content was generally low and particularly poor at position 101 for both B-1a (42%) and CLL (35%) sequences (Figure supplement 6A,D). Collectively, these results demonstrated a progressively lower gradient of Y101 frequencies associated with FCRL6 expression and tissue of origin compared to B-1a and CLL cells (Figure supplement 6E). We concluded that the emerging μHC repertoire in FCRL6^+^ pro B cells, which was inefficient, less diverse, and autoreactive, would disfavor pre-BCR formation and positive selection.

### Higher SLC, V_H_11 and pre-BCR formation by FCRL6^+^ progenitors

Given the unconventional CDR-H3 features in FCRL6^+^ pro B cells and the potential of alternative selection processes (Wong et al., 2017), we focused on the pre-BCR. By RNA-seq, transcripts for the SLC encoding genes *Igll1* and *VpreB1*, were upregulated, whereas *Ikzf1* and *Ikzf3,* which encode factors (Ikaros and Aiolos) that repress SLC genes (Thompson et al., 2007; Ma et al., 2010), were downregulated (Figure 3D - Figure supplement 3A, 4C). Intracellular and surface staining validated these relationships, which were variably more pronounced in the FL than BM (Figure 6A). Aiolos also induces exit from the cell cycle to promote LC rearrangement (Thompson et al., 2007; Ma et al., 2010). Recent studies proposed that premature LC rearrangement by FL pro B cells may obviate pre-BCR formation, leading to direct BCR generation of potentially autoreactive rearrangements, including V_H_12 (Wong et al., 2017). Notably, *Igkc* transcripts were among downregulated overlapping DEGs (Figure 3D). Accordingly, the frequencies of surface κLC were three-fold higher among μHC^+^ FL FCRL6^-^ pro B cells (Figure 6B).

**Figure 6.**
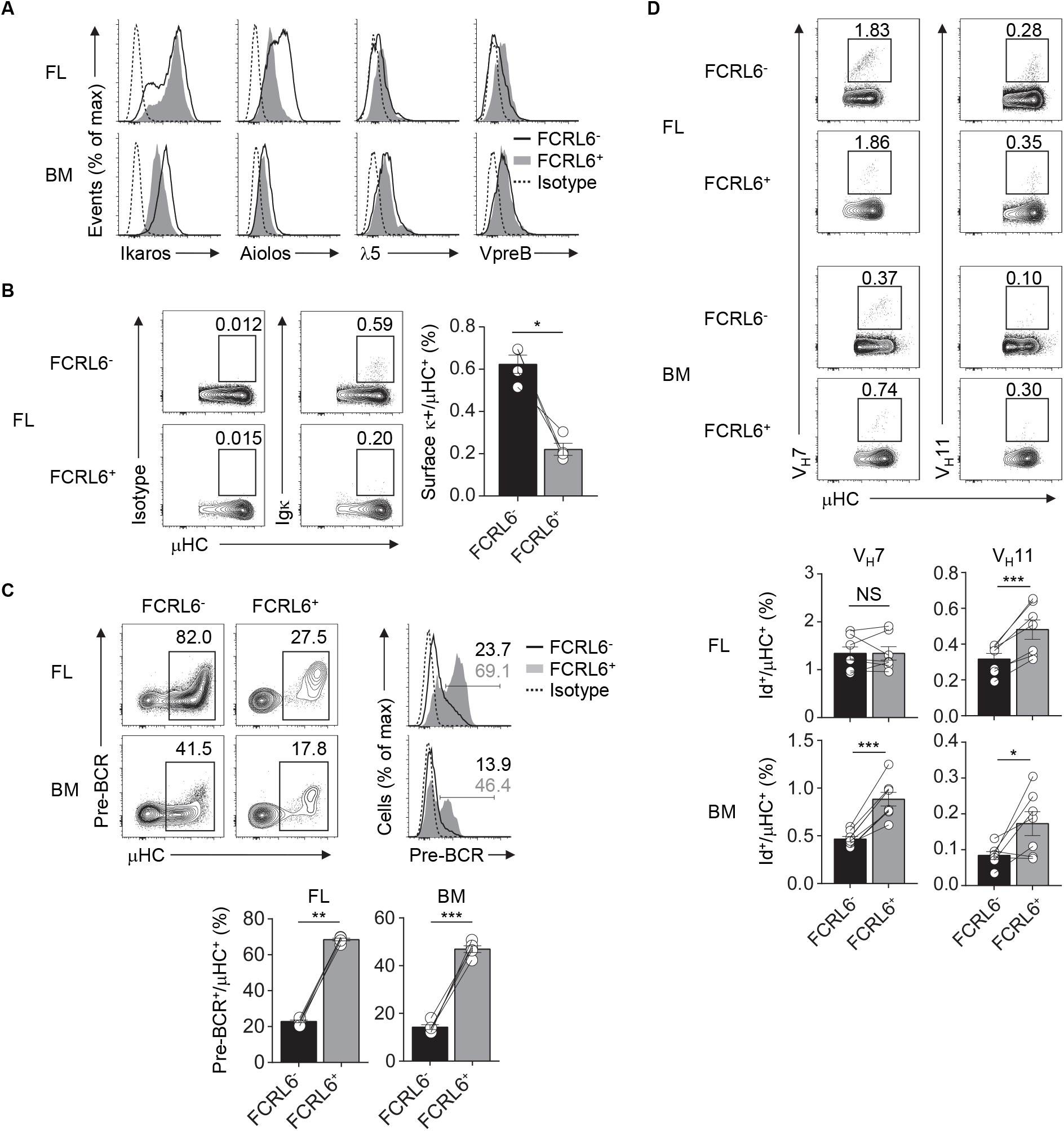
FCRL6 segregates pre-BCR dependent and independent selection and innate-like Ab generation. (**A**) Intracellular and surface staining of transcription factors and SLC components by flow cytometry from FL and BM FCRL6^+^ and FCRL6^-^ pro B cell subsets. (**B**) Flow cytometry analysis of surface κLC expression by μHC^+^ FL pro B cells subsets. Frequencies are indicated adjacent to gated populations. (**C**) Intracellular quantitation of pre-BCR formation by μHC^+^ FL and BM pro B cells. (**D**) Intracellular detection of nascent V_H_7 and V_H_11 within the μHC pool by staining with anti-Id mAbs. Each symbol (**B-D**) represents an individual mouse. Small horizontal lines (**B**-**D**) indicate the mean (± s.e.m.). NS, not significant; \**P* < 0.05, \*\**P* < 0.01 and \*\*\**P* < 0.001 as determined by paired Student’s *t*-test. Data are from 3-5 individual mice and are representative of at least two independent experiments (**A**-**B**), three experiments (**C**), or two (FL) and three (BM) experiments (**D**).

We then examined pre-BCR formation. By co-staining for μHC, we detected 2-3 fold higher pre-BCR levels in FCRL6^+^ pro cells (Figure 6C). Given their unfavorable CDR-H3/Y101 biochemical features, this finding was unexpected. Because *Ighv11* rearrangements were among the most productive in FL FCRL6^+^ cells (Figure supplement 5B,C), we employed anti-idiotype (Id) mAbs to detect the fraction of V_H_11 or V_H_7 μHCs that formed BCRs with emerging PtC or PC reactivity. Although V_H_11 detection was generally rare within the μHC pool (Figure 6D), frequencies were indeed higher among FCRL6^+^ cells. V_H_12 staining was extremely rare among progenitors (data not shown). V_H_7 was more readily detected and while it did not markedly differ between FL subsets, its expression by BM FCRL6^+^ pro B cells was higher. Thus, associated with their disadvantageous CDR-H3 composition, FCRL6^+^ progenitors exhibited higher SLC, V_H_11, and pre-BCR formation, but lower κLC production. These findings suggested the existence of pre-BCR dependent and independent selection processes that varied according to FCRL6 status.

### Myc induction and V_H_11 B-1a development are pre-BCR-dependent

Although FCRL6^+^ cells were mitotically repressed (Figure 3C-F), Myc stimulates early B cell development and proliferation at the pro to pre B cell transition, is repressed by Ikaros and Aiolos, is critical for both B-1 and B-2 lineage development, and is induced in CLL (Habib et al., 2007; Ma et al., 2010; Hayakawa et al., 2016). *Myc* transcripts were upregulated in FL FCRL6^+^ cells by RNA-seq and RQ-PCR (Figure supplement 7A). Furthermore, we confirmed that intracellular Myc expression was elevated in splenic WT B-1a cells and CLL expansions in Eμ-TCL1 Tg mice (Figure supplement 7B). We then investigated Myc as a function of μHC status. While Myc was not appreciably induced in μHC^-^ progenitors, it was upregulated by μHC^+^ FCRL6^+^ B-1P and pro B cells (Figure supplement 7C). These findings were common to both tissues, but were more robust in the FL. To determine if Myc activation was pre-BCR dependent, we analyzed *Igll1* (λ5)^-/-^ mice. These results confirmed that λ5, and thus pre-BCR assembly, was required for Myc activation by FCRL6^+^ μHC^+^ pro B cells (Figure 7A,B). Myc also amplifies calcium signaling in early B cells to promote NFAT activation (Habib et al., 2007), which is required for B-1a development and the generation of PtC-specific reactivity (Berland and Wortis, 2003). NFAT2 was generally higher in FL FCRL6^+^ cells, but levels peaked in FCRL6^+^ μHC^+^ cells (Figure supplement 7D). In the BM, NFAT2 levels were globally lower and differences according to FCRL6 were not as pronounced. However, in contrast to Myc, NFAT2 induction in the FL and BM was independent of pre-BCR formation (Figure 7C-D). Finally, we considered the influence of the pre-BCR on B-1a cell V_H_11 repertoire development. In the C57BL/6 strain, V_H_12 usage between WT and *Igll1* (λ5)^-/-^ B-1a cells did not differ (Wong et al., 2017). We confirmed these findings for V_H_12, but found that V_H_11 reactivity was dramatically lower in both frequency and absolute number in λ5-deficient mice (Figure 7E-F - Figure supplement 7E). In summary, these findings indicate that Myc-induction in FCRL6^+^ progenitor B cells and the development of V_H_11^+^ B-1a cells is pre-BCR dependent.

**Figure 7.**
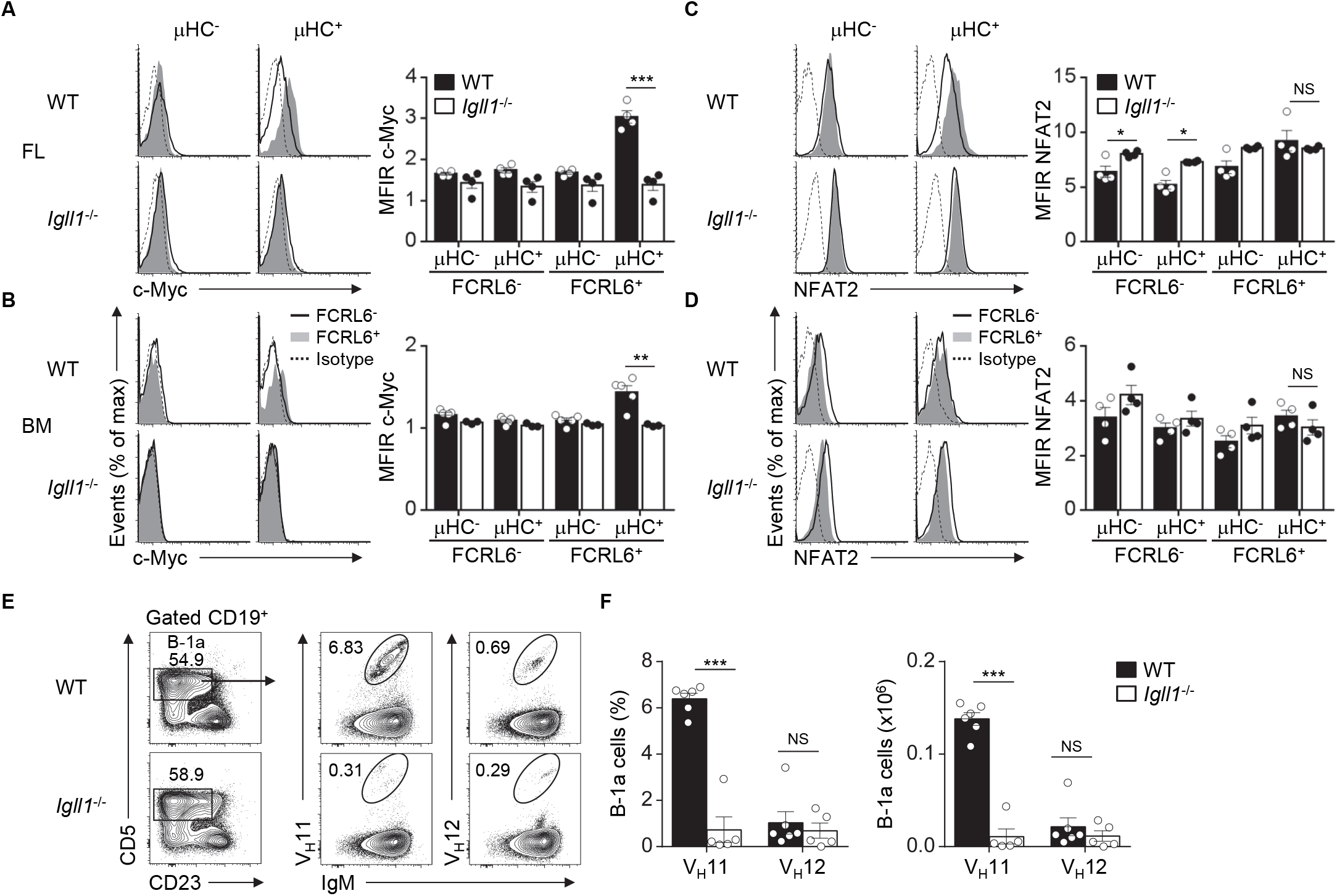
Myc induction by FCRL6^+^ progenitors and V_H_11^+^ B-1a cell development are pre-BCR dependent. (**A**-**B**) Flow cytometry analysis of FL (**A**) and BM (**B**) pro B cells segregated by FCRL6 expression from WT BALB/c and *Igll1*^-/-^ mice intracellularly stained for c-Myc and μHC. (**C**-**D**) Flow cytometry analysis of pro B cells as in **A** and **B** intracellularly stained for NFAT2 and μHC. Data are shown as MFI ratio (MFIR) (test Ab MFI/control Ab MFI). (**E**) Distribution of V_H_11^+^ and V_H_12^+^ B-1a cells from the peritoneal cavities of WT and *Igll1*^-/-^ mice by surface staining with anti-Id mAbs. Numbers in the contour plots indicate frequencies of gated populations from representative mice. (**F**) Frequencies and absolute numbers of V_H_11-Id^+^ and V_H_12-Id^+^ B-1a cells. Each symbol represents an individual mouse. Small horizontal lines (**A**-**D**) indicate the mean (± s.e.m.). NS, not significant; \**P* < 0.05, \*\**P* < 0.01 and \*\*\**P* < 0.001 as determined by paired (**A**-**D**) or unpaired Student’s *t*-test (**E**). Data are from 3-5 mice per strain and are representative of at least two independent experiments (**A**-**F**).

### FCRL6^+^ progenitors exhibit enhanced CXCR5 migration and TSLP differentiation

Resemblance of the FCRL6^+^ pro B cell μHC repertoire and transcriptome to innate-like B cells was further underscored by the induction of gene programs encoding components of innate defense, including Toll-like (*Tlr1*, *Tlr2*, *Tlr4*, *Tlr5*), DNA (*Aim2*, *Ddx58*), and NOD-like (*Nlrc3*) pathogen recognition receptors, as well as cytokine production (*Maf*, *Stat4*, *Ccl3*, *Ccl4*) (Figure 3C,D - Figure supplement 3A,C). Migration and adhesion programs were also upregulated. DEGs encoding the LFA-1 (*Itgb2*) and VLA-4 (*Itga4, Itgb1*) integrins, which direct migration and adhesion to follicular dendritic (FDC) and endothelial cells (Arana et al., 2008), were elevated in FL and BM FCRL6^+^ cells (Figure supplement 8A). CD69 and *Slamf1*/CD150 were also among overlapping DEGs with higher expression by FCRL6^+^ cells. In contrast, we noted that *Ccr7*, which encodes a chemokine that mediates trafficking to T-cell enriched zones (Forster et al., 1999), was downregulated in FL FCRL6^+^ cells and elevated on FL FCRL6^-^ pro B cells (Figure 8A and Figure supplement 3A). These findings indicated different migration properties for these two FL subsets.

**Figure 8.**
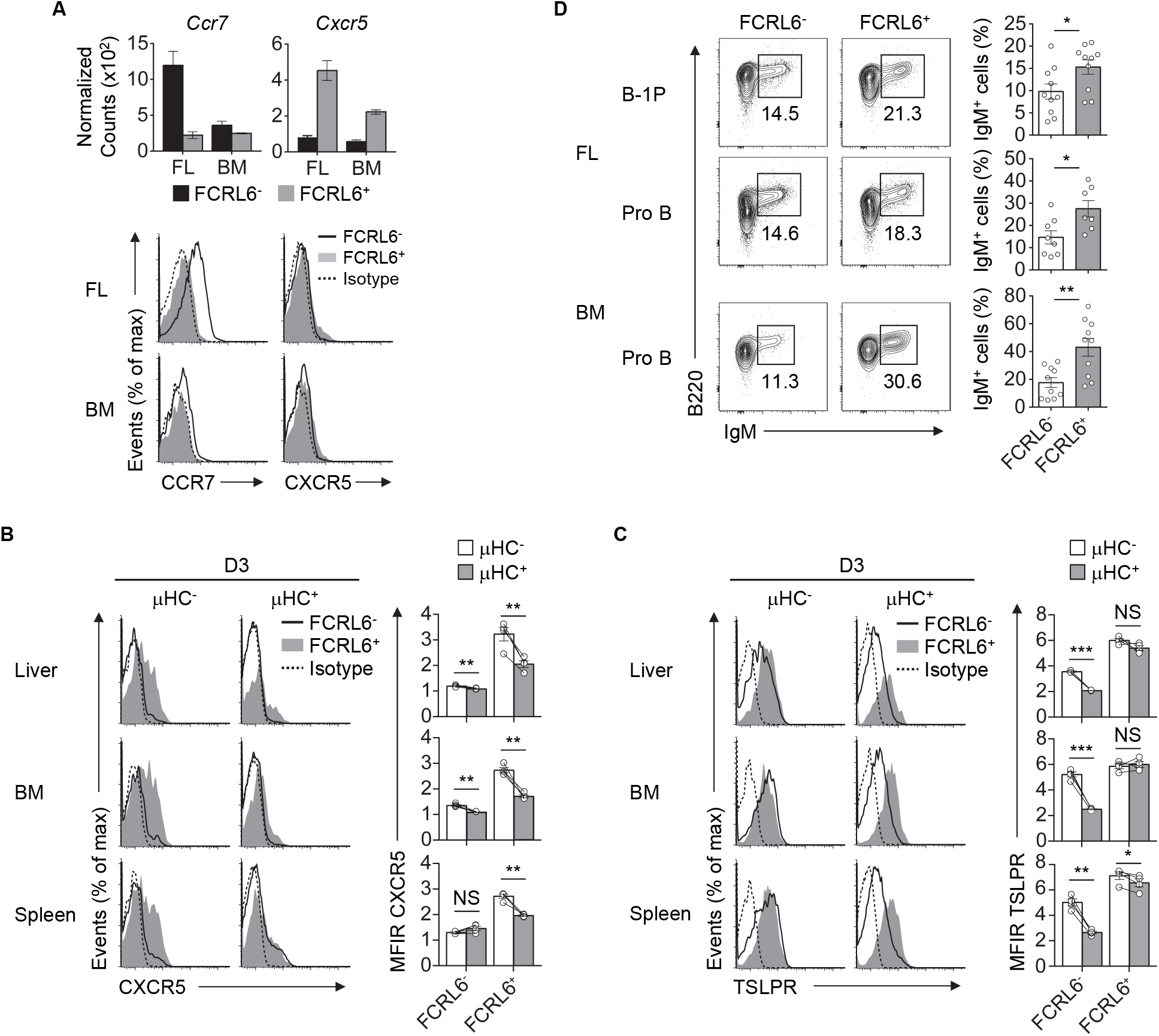
Migration and differentiation of FCRL6^+^ pro B cells is linked to μHC status. (**A**) Normalized counts from RNA-seq (above) and surface staining by flow cytometry analysis (below) of FL and BM pro B cell subsets. Flow cytometry analysis of surface CXCR5 (**B**) or TSLPR (**C**) and intracellular μHC expression in indicated tissues from day 3 (D3) neonatal mice. Data are shown as MFI ratio (MFIR). (**D**) Differentiation of sorted FL and BM FCRL6^+^ and FCRL6^-^ progenitor B cell subsets cultured in transwell plates over 7-8 days with the S17 stromal cell line and cytokines according to Montecino-Rodriguez et al., 2006. Small horizontal lines (**B-D**) indicate the mean (± s.e.m.). \**P* < 0.05, \*\**P* < 0.01 and \*\*\**P* < 0.001 as determined by paired Student’s *t*-test. Data are representative of two independent experiments (**A**), pooled tissues from one of two independent experiments (**B-C**), and cumulative data from 5-6 individual experiments (**D**).

B-1a cells express CXCR5 and home to the PeC and body cavities in a CXCL13-dependent fashion (Ansel et al., 2002). Although *Cxcr5* was among upregulated overlapping DEGs, it was not detected on FCRL6^+^ pro B cells (Figure 3D and 8A). However, CXCR5^+^ transitional B cells entering the spleen 18 hours before birth acquire postnatal CXCL13 responsiveness to initiate white pulp formation (Neely and Flajnik, 2015). Thus, we postulated that CXCR5 might be dynamically induced to direct migration after parturition. We therefore examined CXCR5 in the liver, BM, and spleen from day 3 and 7 neonates. CXCR5 surface density and cell frequency was uniformly higher on FCRL6^+^ progenitors among day 3 tissues (Figure supplement 8B). Although FCRL6^-^ pro B cells had lower CXCR5, it escalated on subpopulations in the spleen > BM > liver. At day 7, largely homogenous CXCR5 expression was still evident for liver FCRL6^+^ pro B cells, but it bifurcated in the BM and was nearly lost in the spleen. Subpopulations of CXCR5^+^ FCRL6^-^ pro B cells were detectable at day 7 in liver and BM, but a distinct subset emerged in the spleen. Given the impact of pre-BCR formation on Myc induction in FCRL6^+^ cells, we examined CXCR5 as a function of μHC status. At day 3, CXCR5 levels on FCRL6^-^ pro B cells were generally low regardless of μHC expression (Figure 8B). In contrast, elevated CXCR5 expression by FCRL6^+^ μHC^-^ cells markedly declined in μHC^+^ cells. These results indicated that CXCR5 and μHC production were linked in FCRL6^+^ cells, and that migrating μHC^-^ cells repress CXCR5 upon becoming μHC^+^.

We then considered the impact of TSLP, a trophic factor capable of driving differentiation of fetal B cell progenitors to surface IgM^+^ cells (Levin et al., 1999). FCRL6^-^ pro B cells demonstrated higher IL-7Rα expression, but TSLPR levels did not markedly differ (Figure 2A and data not shown). TSLPR expression by FCRL6^+^ and FCRL6^-^ B cell progenitors was similar among day 3 and 7 postnatal tissues (Figure supplement 8C). However, with respect to μHC status, μHC^+^ FCRL6^-^ cells strongly downmodulated the TSLPR (Figure 8C). In contrast, sustained TSLPR expression by μHC^+^ FCRL6^+^ cells indicated that responses to this cytokine differed between these subpopulations. To investigate this, we first sorted BM FCRL6^+^ and FCRL6^-^ pro B cells and performed *in vitro* growth assays in the presence of TSLP. Survival of both subsets increased in a dose-dependent manner; however, FCRL6^+^ pro B cells exhibited greater TSLP responsivity (Figure supplement 8D). We next examined differentiation by adopting an *in vitro* co-culture system established by the Dorshkind group that includes TSLP (Montecino-Rodriguez et al., 2006). In initial studies, we detected few IgM^+^ B cells at day 4 and by day 12, most cells were dead (data not shown). At day 8, however, sorted FCRL6^+^ FL B-1P and FL and BM pro B cells were consistently more proficient than FCRL6^-^ cells in differentiating to IgM^+^ cells (Figure 8D). Thus, FCRL6^+^ progenitors exhibited preferential CXCR5- and TSLPR-mediated homing and differentiation potential.

## Discussion

Here FCRL6 defined a subset of B cell progenitors that is conserved throughout ontogeny. FCRL6 expression delineated two pro B cell subpopulations with marked differences in their molecular signatures, biology, and emerging μHC repertoires. Their distinct rearrangement activities, migration potential, and differentiation pathways culminated in pre-BCR dependent and independent modes of selection. This disparity further correlated with the developmental potential of two B-1a stereotypic clones that give rise to natural Abs.

The origin of the B-1 lineage has been the subject of considerable discussion. Despite progress in identifying B-1 progenitors (Montecino-Rodriguez et al., 2006), a shortfall in the support of the ‘lineage’ theory has been a lack of definitive regulatory factors that uniquely drive B-1 development. Identification of the Let7-Lin28b-Arid3a axis has provided supportive evidence for a developmental switch (Montecino-Rodriguez and Dorshkind, 2012; Yuan et al., 2012; Zhou et al., 2015; Kristiansen et al., 2016). However, *in vitro* provision of these elements did not recapitulate expected biases of natural Ab repertoires enriched with the PtC-reactive V_H_11 and V_H_12 families (Zhou et al., 2015). Thus, while this circuit may differentially impact features of B-1 versus B-2 differentiation that predominate in fetal and adult life (*i.e.*, phenotype, tissue localization), some programmatic regulation (*i.e.*, repertoire formation and selection) may be independent of this governance.

Perhaps the strongest evidence for a B-1a specific regulator is *Bhlhe41.* Mice deficient in this factor have dramatic reductions in B-1a cells from the transitional stage of maturation, altered Ab repertoires, and a near complete loss of V_H_12 BCRs (Kreslavsky et al., 2017). However, V_H_11 usage in the absence of Bhlhe41 appears intact. Notably, these studies differ from ours by the use of C57BL/6 background mice. However, findings that *Fcrl6* and *Bhlhe41* were the two most differentially expressed genes between FL and BM subsets in our study strongly imply disparate regulatory outcomes related to FCRL6 expression. This is further highlighted by: (i) the increased efficiency of V_H_11 μHC generation and pre-BCR formation by FCRL6^+^ cells, and (ii) the marked loss of V_H_11, but not V_H_12 B-1a development in *Igll1*^-/-^ mice. Hence, these findings indicate that differences in Ab generation and selection are hard wired. The regulatory factors responsible may relate to nervous system-related DEGs identified by the GO pathway analysis. *Tcf7*/*Lef1* genes, which regulate Wnt/β-catenin signaling, self-renewal, and early B cell development, are attractive candidates (Reya et al., 2000). *Bcl11a*, *Foxo1*, and *Klf3* were also among DEGs, but how these factors influence *Igh* repertoire formation with respect to the newfound heterogeneity disclosed here, remains unclear.

The B-1a BCR repertoire is understood to be driven by Ag-dependent positive selection (Hayakawa et al., 1999; Graf et al., 2019). Here, we identified pro B cell subset-specific *Igh* repertoire skewing that became evident from the onset of recombination through pre-BCR selection. Despite elevated *Rag1*/*2* expression and cell cycle enrichment in G0/G1, DNA repair was globally repressed in FCRL6^+^ progenitors. Furthermore, clear differences in proximal versus distal *Ighv* locus accessibility, V(D)J productivity, CDR-H3 composition, diversity, and μHC expression, illustrated their developmental divergence. These findings imply that FCRL6^+^ cells serve an ancient, but fundamental role as specialized producers of natural Abs. However, this capacity may not be exclusive, as we detected V_H_11-Id^+^ μHC in FCRL6^-^ progenitors. Evidently, mechanistic restrictions inherent to FCRL6^+^ cells serve to restrain Ab diversification by sculpting rearrangements that encode charged or hydrophobic CDR-H3 features that are enriched in innate-like MZ and B-1a B cells (Khass et al., 2018). Because FCRL6 expression is lost following positive checkpoint selection, we did not examine μHC sequences after this point in differentiation. Thus, distinguishing FCRL6^+^ from FCRL6^-^ progenitors following this development stage will require lineage tracing.

The increased efficiency of pre-BCR formation, Myc induction, and consequences of μHC expression (*e.g.*, migration, differentiation), indicate positive selection and differentiation of FCRL6^+^ cells along a distinct developmental path. However, the promotion of μHCs harboring CDR-H3s with disadvantageous composition through the pre-BCR checkpoint, contrasts with the notion that such autoreactive features trigger negative selection to maintain central tolerance (Keenan et al., 2008). This unexpected result, in light of the importance of the Y101 residue (Bankovich et al., 2007; Khass et al., 2016), might suggest that non-CDR-H3 interactions contribute. It is intriguing that autonomous BCR signaling in CLL involves an FR2 epitope (Duhren-von Minden et al., 2012). Perhaps, nascent μHC/SLC interactions of low affinity are compensated by altered stoichiometry. Ikaros and Aiolos repression correlated with VpreB/λ5 upregulation in FCRL6^+^ cells. Emerging CDR-H3s with unfavorable H-bond potential could become saturated by excess SLC that precipitates pre-BCR formation. Evidence that premature κLC rearrangement obviates this checkpoint (Wong et al., 2017), was also verified here, but appears to differ according to FCRL6 expression. Thus, at least two pathways appear to promote the development of μHC^+^ innate-like progenitors. Furthermore, these modes of pre-BCR dependent and independent passage correlate with V_H_11 and V_H_12 B-1a development and Myc induction.

Progenitors marked by FCRL6 exhibited a strong transcriptomic, phenotypic, and *Ighv* repertoire resemblance to CLL cells. In addition to their emerging autoreactivity, CLL-related transcription factors (LEF-1, NFAT2), innate-defense components (TLRs), chemokines/cytokines (Ccl3, Ccl4), migration elements (CXCR5), and Myc expression were also shared. Hayakawa found that both BCR autoreactivity and B-1a physiology were required for CLL leukemogenesis in mouse models (Hayakawa et al., 2016). Our data demonstrate that FCRL6^+^ progenitors exhibit both these properties and have the potential to home to sites where chronic stimulation and innate inflammation could fuel their transformation over time.

The interplay between developing B cell progenitors and the microenvironment into which they are born and ultimately home to during ontogeny is dynamic and physiologically complex. Based on the marked shift in biology triggered by μHC expression and pre-BCR assembly on Myc, CXCR5, and TSLPR, it appears that perinatal FCRL6^+^ cells are poised for migration even prior to μHC expression. These findings are in line with those by Wen et al. who demonstrated the association of CXCR5^hi^-expressing V_H_11^+^ B-1a cells with white pulp FDCs in the neonatal spleen (Wen et al., 2005). In this fetal microenvironment, FCRL6^+^ cells could contribute to both architectural changes of developing secondary lymphoid tissues and innate-like humoral defense prior to parturition.

The function of FCRL6, its ligand(s), and regulatory influence as a defining marker is unknown. Ig binding studies with transductants and by surface plasmon resonance did not reveal evidence of Ab binding (data not shown). The lack of a canonical tyrosine-based cytoplasmic motif suggests that FCRL6 likely recruits different intracellular effectors than other FCRL family members. A hurdle in modeling its role is a lack of cell lines that express it and thus *in vivo* systems will be required to define its physiological functions.

In summary, we have taken an unbiased approach to investigating a defining marker of early B lineage differentiation. This work has yielded important insight into the developmental heterogeneity, origins, and selection of innate-like B cells. The frequencies of FCRL6^+^ cells reflect the B-1 potential of fetal and adult tissues and their biased polyreactive repertoires, genetic programs, and migratory features are shared with MZ, B-1a, and CLL cells. These findings introduce a model that integrates mechanisms associated with both the ‘lineage’ and ‘selection’ theories of B-1 development at the beginning stages of innate-like B cell differentiation.

## Materials and Methods

### Mice

BALB/cJ and C57BL/6J, as well as μMT and *Rag2*^-/-^ mice on the C57BL/6 background, were purchased from Jackson Laboratories and bred and maintained in animal facilities at the University of Alabama at Birmingham (UAB). *Dntt*^-/-^ (Tdt), *Igll1*^-/-^ (λ5), and *Rag1*^-/-^ mice on the BALB/c background were generously provided by Dr. Harry Schroeder Jr. at UAB. Eμ-TCL1 Tg mice were kindly provided by Dr. Carlo Croce at Ohio State University. Unless otherwise specified, 8-12 week-old female mice were used for these studies. Embryonic fetal livers (FL) were obtained from timed pregnancies. Vaginal plug formation after mating was counted as day 0. All studies and procedures were approved by the UAB Institutional Animal Care and Use Committee (IACUC).

### Quantitative PCR

Total RNA was extracted from mouse tissues and single cell suspensions using RNeasy Plus kits (Qiagen). cDNAs were generated using SuperScript II (Invitrogen). cDNAs were mixed with primer pairs and amplified using SYBR green master mix and the 7900HT Fast real-time PCR system (Applied Biosystems). Iμ, μ0, and *Myc* primers have been published (Kuzin et al., 2008; Featherstone et al., 2010; Sumner et al., 2017). *Fcrl6* qPCR primers were designed to hybridize with the first extracellular domain using primer express software (Applied Biosystem). Samples were normalized to *Polr2a (RNA polymerase II*) expression (Radonic et al., 2004). Primer sequences were as follows:

*Fcrl6* F: 5’-CATGCTGCTCTGGATGGTTCT-3’
*Fcrl6* R: 5’-AGCTCAGGATTTGGGAACAACTC-3’
Iμ F: 5’-GGATACGCAGAAGGAAGGC-3’
Iμ R: 5’-GGTCATTACTGTGGCTGGAGAG-3’
μ0 F: 5’-TGCAGGTTCCTCTCTCGTTTCCTT-3’
μ0 R: 5’-TGGGCCCATCTGTAGGATGGTAAT-3’
*Myc* F: 5’-AACAGGAACTATGACCTCG-3’
*Myc* R: 5’-AGCAGCTCGAATTTCTTC-3’
*Polr2a* F: 5’-GACTCACAAACTGGCTGACAT-3’
*Polr2a* R: 5’-TACATCTTCTGCTATGACATGGG-3’

### Generation of anti-mouse FCRL6 antibodies (Abs)

Rat anti-mouse FCRL6-specific mAbs, 1C3 (IgG1κ) and 3C1 (IgG2aκ), were generated using *Escherichia coli*-derived His-tagged recombinant protein comprised of the two extracellular Ig domains of mouse FCRL6. Respective cDNA regions were PCR amplified and cloned into the pET24b vector (Novagen) for bacterial expression, as previously described (Alder et al., 2005). Fisher rats (Jackson Laboratory) were immunized at 3 to 4 day intervals over a 3 week period and popliteal nodes were fused with the mouse plasmacytoma Ag8.653 and plated for selection in 96-well plates (Won et al., 2006). At 10–14 days after fusion, hybridoma clones were screened for specificity and cross-reactivity by staining hemagglutinin (HA)-tagged FCRL1, FCRL5 (C57BL/6 and BALB/c allelles), and FCRL6 TM transductants generated as previously (Won et al., 2006). Rabbit anti-FCRL6 polyclonal Abs were generated by hyperimmunizing New Zealand White rabbits (Charles River Laboratories) with *Escherichia coli*-derived His-tagged recombinant protein.

### Immunoprecipitation and Western blotting

To analyze the molecular nature of FCRL6, HA-tagged BW5147 FCRL6 retroviral transductants (1 x 10^7^) were lysed in 1% NP-40 lysis buffer. Whole cell lysate proteins were quantitated using the BCA reagent (Pierce) and incubated at 4°C for 30 min with rat anti-mouse FCRL6 (3C1) or an isotype-matched control (rat IgG2aκ) mAb, followed by the addition of 30 μl of a 50% slurry of protein G beads (GE Healthcare), and incubation overnight at 4°C. The beads were washed five times with 1 ml of lysis buffer to reduce non-specific binding, resuspended in an equal volume of SDS sample buffer, and boiled. Proteins were resolved by SDS-PAGE and transferred to PDVF membranes (Millipore) before immunoblotting with anti-HA (12CA5, Roche) or rabbit anti-mouse FCRL6 polyclonal Abs, followed by rabbit anti-mouse or goat anti-rabbit HRP (Southern Biotech). Bound Abs were visualized using the ECL reagent (GE Healthcare) and detected by Biomax XAR film (Kodak).

### Flow cytometry and cell sorting

Spleen, FL, and bone marrow (BM) cells were prepared as single cell suspensions after red blood cell lysis with ACK lysing buffer (Gibco). Peritoneal cavity cells were prepared by lavaging with complete 10% FCS RPMI-1640 medium. FL were obtained from timed pregnancies. Neonatal livers, spleens, and BM were harvested at specific days after birth. Cells were stained with antigen–specific or isotype-matched control Abs (key resources table) after blocking with unlabeled anti-CD16/32 for 5 min.

AF647 and biotin-conjugated anti-mouse FCRL6 (1C3 and 3C1) were generated and labeled by our laboratory (Invitrogen and Thermo Scientific). Anti-V_H_11 (3H7 (Rowley et al., 2007); a gift from K. Hayakawa), anti-V_H_12 (5C5 (Arnold et al., 1994); a gift from K. Rajewsky), and anti-V_H_7 (TC68 (Desaymard et al., 1984), a gift from J. Kearney) were biotinylated in our laboratory (Thermo). Cells were analyzed using FACSCalibur (BD) or LSRII (BD) flow cytometry instruments and plotted with FlowJo software (Treestar).

For cell sorting, leg bones were pooled from 5-10 adult male and female mice and 7-10 FL were pooled from E18 fetuses isolated from the wombs of pregnant mothers according to the timing of vaginal plug formation. Single cell suspensions from the BM of BALB/cJ mice were prepared by flushing the femur and tibia bones with a 29 G needle syringe. Single suspensions of FL cells were prepared by crushing the tissue with a 1 ml syringe plunger and passage through a 40 μm strainer (BD). Single cell suspensions were treated with ACK lysing buffer and re-suspended with PBS containing 2% FCS. After Fc blockade with unlabeled anti-CD16/32, cells were stained with fluorochrome-labeled mAbs (FITC-CD43, BV421-CD19, PE-B220, PECy7-AA4.1, and Alexa647 FCRL6 (1C3)), as well as biotin-labeled mAbs (CD3 (145-2C11), Mac1 (M1/70), Gr1 (RB6-8C5), Ly6C (HK1.4), Ter119, and IgM (RMM-1) to exclude T, myeloid, and erythrocyte lineage cells and immature/mature B cells, and counterstained with SA-BV570. FCRL6^+^/FCRL6^-^ pro B (AA4.1^+^Lin^-^CD43^+^B220^+^CD19^-^IgM^-^) cells from the FL and BM were sorted using a FACSAria II sorter (BD).

### *In vivo* proliferation and cell cycle analysis

Single cells were prepared from the FL and BM of BALB/cJ mice at 24 hrs after E17 pregnant dams or adult mice were injected i.p. with BrdU (twice at 12 hr intervals). Single cell suspensions were stained with anti-mouse CD93, CD43, CD19, B220, IgM, and FCRL6 (1C3) for surface detection, then fixed and permeabilized, treated with DNase I, and stained with anti-BrdU and 7AAD to examine proliferation and cell cycle status. Stained cells were analyzed by FACS with an LSRII instrument and profiles were plotted with FlowJo software.

### Intracellular staining

Single cells from FL and BM were stained for CD93, CD43, CD19, B220, IgM, and FCRL6 (3C1), and fixed with Cytofix (BD) for 15 min on ice, then permeabilized with Foxp3 Fixation/Permeabilization buffer (eBio) for 30 min on ice, and stained for either Ki-67, c-Myc, NFAT2, Ikaros, or Aiolos, along with F(ab’)_2_ goat anti-mouse IgM for 1 hr at room temperature. Cells were examined using an LSRII cytometer and plotted with FlowJo software.

### Phospho-flow analysis

FACS sorted FCRL6^+^ and FCRL6^-^ pro B (CD43^+^CD19^+^B220^hi^IgM^-^) cells from adult BM were treated with the phosphatase inhibitor pervanadate (NaVO_4_) for 10 min. Stimulated cells were fixed with prewarmed Phosflow Lyse/Fix buffer (BD) at 37°C for 10 min and permeablized with Phosflow Perm Buffer III (BD). After Fc blockade (CD16/32), cells were stained with anti-phospho ERK pT202/pY204, STAT5 Y694, or isotype control mAbs (BD) for 30 min at room temperature. Phosphorylation was analyzed using a FACSCalibur flow cytometer (BD) and plotted with FlowJo software. The fold induction change in phosphorylation for FCRL6^-^ and FCRL6^+^ pro B cells was calculated by comparing the MFI ratios of FCRL6^-^/FCRL6^+^ pro B cells with and without stimulation.

### Pre-BCR and intracellular IgM staining

FCRL6^+^ and FCRL6^-^ pro B cells (CD93^+^CD43^+^CD19^+^B220^hi^IgM^-^) stained for cell surface markers, were fixed with Cytofix/Cytoperm buffer (BD) for 20 min on ice and stained with anti-pre-BCR (SL156) and/or F(ab’)_2_ goat anti-mouse IgM for 1 hr at room temperature. Cells were analyzed using an LSRII instrument and plotted with FlowJo software.

### Apoptosis assays

Single cells from the FL and BM were stained for cell surface markers then washed twice in Annexin V binding buffer, followed by staining with anti-Annexin V for 15 min at room temperature. After incubation with the Annexin V binding buffer, cells were analyzed using an LSRII instrument and plotted with FlowJo software.

### *In vitro* proliferation assays

FACS-sorted FCRL6^+^ and FCRL6^-^ pro B cells (AA4.1^+^ CD43^+^CD19^+^B220^hi^Lin^-^IgM^-^) from BM were washed twice with complete RPMI 1640 media containing 10% FCS. Pro B cells (10^3^ cells per well) were loaded into round-bottom 96-well plates in triplicate, in the presence or absence of different concentrations of TSLP (0-100 ng/ml). Cells were incubated at 37°C in a CO_2_ incubator for 4 days, total cell numbers from each well were counted, and live cells were enumerated by flow cytometry analysis using a FACScalibur instrument with dead cell exclusion by PI.

### Progenitor differentiation assays

Stromal cell-dependent B cell progenitor differentiation assays were previously described by Montecino-Rodrigez et al. and performed as follows (Montecino-Rodriguez and Dorshkind, 2006; Montecino-Rodriguez et al., 2006). S17 stromal cells were freshly thawed and grown in complete RPMI with 10% FCS for 7-10 days. S17 cells were detached with 0.25% Trypsin-EDTA solution (Gibco), filtered with a 70 μm cell strainer (BD), and 2.5 x 10^4^ cells were added to 6-well Biocoat Transwell insert plates (BD) for growth on inserts overnight. FACS-sorted FL or BM B-1P and pro B cells (0.5-1 x 10^4^ cells/well), gated as in Figure 1B, were added to the bottom chamber of the Transwell plate, separated from S17 stromal cells in the upper chamber, and cultured in αMEM media (Gibco) with 5% FCS and a cytokine cocktail including TSLP (10 ng/ml), IL-3 (5 ng/ml), IL-6 (10 ng/ml), SCF (10 ng/ml), and Flt3L (10 ng/ml). At day 4, 25% of the upper and lower culture media was replaced with fresh cytokine-containing media. At day 8, the cells were harvested and washed with PBS containing 2% FCS. Viability, proliferation, and differentiation into immature B220^+^IgM^+^ B cells were examined with an LSRII cytometer (BD) and analyzed with Flow-Jo software.

### *Ighv* repertoire sequencing and analysis

RNA from FCRL6^+^ to FCRL6^-^ pro B cells sorted as in Figure 1B, from pooled E18 FL (*n* = 7) or adult BM samples (*n* = 8) from BALB/c mice, was isolated using TRIzol (Invitrogen) reagent and used to generate cDNA with the High-Capacity cDNA synthesis kit (Applied Biosystems). cDNAs were used as a template for PCR using the KOD high fidelity polymerase (Novagen) and a mixture of 8 V_H_ family and IgM-specific primers (listed below) containing 5’ adaptors to allow barcoding by the Illlumina TruSeq multiplex pcr kit (Illumina). Resulting PCR products were subjected to 10 cycles of PCR using Illumina TruSeq indexing primers. The resulting amplicons were purified using AMPure (Beckman), submitted to the UAB Heflin Genomics core, and sequenced using Illumina’s MiSeq Reagent Kit v3 2X300 kit in a Miseq next generation sequencer. The resulting FastQ files were filtered for quality (>Q30) and assembled using the vsearch suit of informatic tools (Rognes et al., 2016). Fasta files containing the full length paired *Igh* sequences were submitted to the IMGT High-VQuest server for *Ighv*, *Ighd*, and *Ighj* identification, as well as *Ighv* junction and other bioinformatic analyses (Brochet et al., 2008). IMGT files were visualized using R software (R Foundation for Statistical Computing) and the following software packages: vegan, plyr, plots, ggplots, ggrepel, ggseqlogo, pbmcapply, seqinr, and stringr. Some rarefaction curves were replicated with VDJtools. Sequences were deposited at NCBI accession: PRJNA547609.

#### *Ighv*-specific primer sequences

An i5 linker (5’-CCC TAC ACG ACG CTC TTC CGA TCT-3’) was added to the 5’ end of the forward V_H_ primers and an i7 linker (5’-ATC TCG TAT GCC GTC TTC TGC TTG-3’) to the 5’ end of the reverse Cμ primer.

VH1a F: 5’-CAGGTGTCCACTCCCAGGTCC-3’
VH1b F: 5’-CAGGTGTCCTCTCTGAGGTCCAG-3’
VH2/4 F: 5’-CAGGTGTCCTATCCCAGGTGCAGC-3’
VH3 F: 5’-GAAGGTATCCTGTCTGATGTGCAGC-3’
VH8 F: 5’-CAGATGTCCTGTCCCAGGTTACTC-3’
VH9/11/12 F: 5’-CAGGTGCCCAAGCACAGATCCAG-3’
VH5/6/7/10/13 F: 5’-CAGGTGTCCAGTGTGAAGTGCAGC-3’
VH14 F: 5’-CAGAGGTTCAGCTGCAGCAGTCTGG-3’
Cμ R: 5’-AGGGGGAAGACATTTGGGAAGGAC-3’

### Sequence analysis of the CDR-H3 regions

CDR-H3 analysis was performed as in Khass et al (Khass et al., 2016). Briefly, the CDR-H3 was defined as the sequence starting immediately after the cysteine at the end of FR3 at position 96 and then through, but not including, the tryptophan that begins FR4. At positions 99 to 103, we calculated the frequency of amino acids for individual B cell subsets and common *Ighv* segments. Logo plots were prepared with the ggseqlogo R package (Wagih, 2017).

### RNA-seq library preparation and analysis

RNA sequencing (RNA-seq) was performed at the UAB Heflin Center for Genomic Science. E18 FL and adult BM FCRL6+/- pro B cells were sorted in duplicate in four independent experiments from BALB/cJ mice by flow cytometry as detailed above and shown in Figure 1B. mRNA was isolated using RNeasy Plus kits (Qiagen) and sequencing was performed on a Illumina NextSeq 500 (paired-end sequencing 2×75bp) according to the manufacturer’s guidelines. Briefly, the quality of the total RNA was assessed using the Agilent 2100 Bioanalyzer (all samples had RIN values of >9.9) followed by two rounds of poly(A) selection and conversion to cDNA. TruSeq library generation kits were constructed by random fragmentation of the polyA mRNA, followed by cDNA production using random primers (Agilent Technologies). The ends of the cDNA were repaired, A-tailed, and adaptors were ligated for flow cell attachment, sequencing, and indexing as per the manufacturer’s instructions (Agilent Technologies). cDNA libraries were quantitated using qPCR in a Roche LightCycler 480 with the Kapa Biosystems kit for library quantitation (Kapa Biosystems) prior to sequencing.

All samples contained a minimum of 32 million reads with an average number of 42.9 million reads across all biological replicates. The FASTQ files were uploaded to the UAB High Performance Computer cluster for bioinformatics analysis with the following custom pipeline built in the Snakemake workflow system (v4.8.0) (Koster and Rahmann, 2012): first, quality and control of the reads were assessed using FastQC, and trimming of the bases with quality scores of less than 20 was performed with Trim_Galore! (v0.4.5). Following trimming, the reads were aligned with STAR (Dobin et al., 2013) (v2.5.2a, during ‘runMode genomeGenerat’, the option ‘sjdbOverhang’ was set to 74) to the Ensembl mouse genome (mm10), which resulted in an average of 80.3% uniquely mapped reads. BAM file indexes were generated with SAMtools (v1.6) and gene-level counts were generated using the function ‘featureCounts’ from the R package, Rsubread (v1.26.1; r-base v3.4.1), with the ‘Mus_musculus.GRCm38.90.gtf’ file from Ensembl. The parameters used in ‘featureCounts’ included: isGTFAnnotationFile=TRUE, useMetaFeatures=TRUE, allowMultiOverlap=FALSE, isPairedEnd=TRUE, requireBothEndsMapped=TRUE, strandSpecific=2, and autosort=TRUE. Additional parameters were kept as default. Logs of reports were summarized and visualized using MultiQC (v1.5) (Ewels et al., 2016).

Count normalization and differential expression analysis were conducted with R software with the DESeq2 package (Love et al., 2014) (v1.16.1). Following count normalization, principal component analysis (PCA) was performed (Figure 3A) and genes were defined as differentially expressed genes (DEGs) if they passed a statistical cutoff containing an adjusted *P*-value <0.05 (Benjamini-Hochberg False Discovery Rate (FDR) method) and if they contained an absolute log_2_ fold change >=1. Functional annotation enrichment analysis was performed in the NIH Database for Annotation, Visualization and Integrated Discovery (DAVID, v6.8) by submitting all DEGs identified. The Benjamini-Hochberg FDR correction was also applied to determine gene ontology (GO) terms with the cutoff of an adjusted *P*-value <0.05.

RNA-seq files are available at the Gene Expression Omnibus under accession number GSE132438. Figures, including the heatmap, scatter plots, and volcano plot, were made with R software using the following packages: ggplot2 (v3.1.0) and ComplexHeatmap (v1.20.0).

### Statistical analysis

Statistical analysis was performed using GraphPad Prism 7 software. A paired Student’s *t*-test was used for comparing two experimental groups (FCRL6^+^ and FCRL6^-^ progenitors) for apoptosis, intracellular and surface staining, RQ-PCR, and differentiation assays. Unpaired Student’s *t*-test was used for comparing cell populations in mutant mouse studies. Statistical analyses used in RNA-seq studies are detailed in the indicated section.

## Acknowledgements

We thank J. Lama, J. Wu, D. Brooke, A. Foksinska, M. Amjad, Z. Zhu, and A. Weinmann for technical assistance; C.M. Croce (Ohio State University) for providing Eμ-TCL1 Tg mice; H. W. Schroeder Jr. for providing *Dntt*^-/-^, *Igll1*^-/-^ and *Rag1*^-/-^ mice; J.F. Kearney for providing μMT mice, S-17 cells, and anti-V_H_7; K. Hayakawa (Fox Chase Cancer Center) for anti-V_H_11, K. Rajewsky (Max Delbrück Center for Molecular Medicine) for anti-V_H_12; and X.J. Yan and N. Chiorazzi (Feinstein Institute for Medical Research) for unpublished mouse CLL sequences. We appreciate the assistance of V. S. Hanumanthu (UAB - Comprehensive Flow Cytometry Core - NIH P30 AR048311 and AI27667) and M. Crowley (UAB - Heflin Center for Genomic Sciences). The authors thank P.D. Burrows, J.F. Kearney, and H.W. Schroeder Jr. for critically reading the manuscript. This work was supported in part by the UAB CLL and Cancer Immunobiology Programs (R.S.D).

---

**Figure supplement 1.**
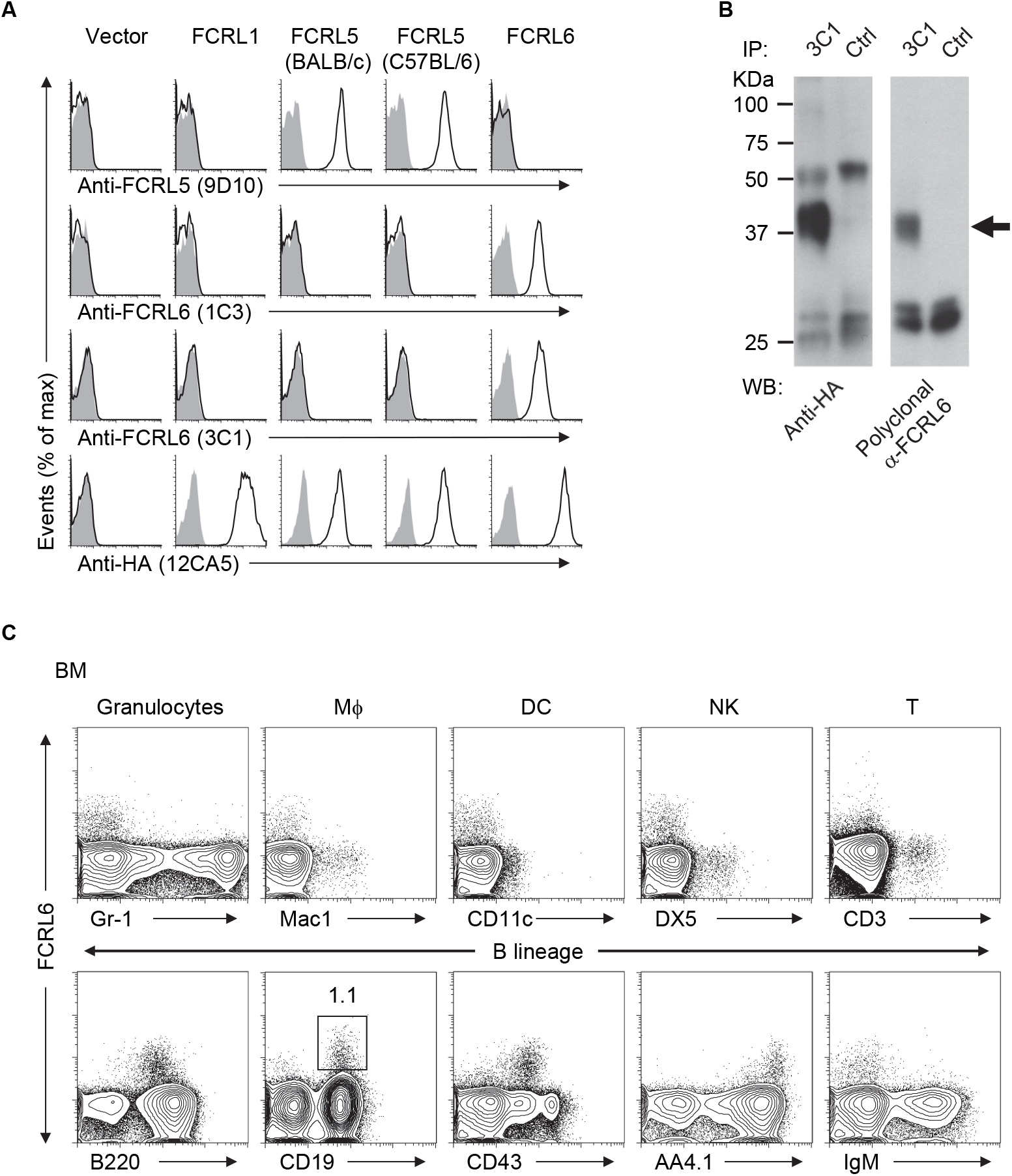
Specificity of FCRL6 monoclonal antibodies (mAb) and early B cell restricted expression. (**A**) Specificity of rat anti-mouse FCRL6 mAbs. *E. coli*-derived recombinant protein of the two FCRL6 Ig-like extracellular domains was injected into Fisher rats and mAbs were generated by standard techniques as previously (Won et al., 2006). Specificities of the 1C3 and 3C1 subclones were tested for cross-reactivity with a panel of BW5147 retroviral transductants expressing HA-tagged mouse FCRLs by staining with the indicated mAbs prior to flow cytometry analysis (black lines) or an isotype-matched control (gray shade). (**B**) Molecular nature of FCRL6 (indicated by the arrow). BW5147 FCRL6 transductants were immunoprecipitated with the 3C1 mAb or an isotype control (rat IgG2aκ) and blotted with an anti-HA (12CA5) mAb or rabbit anti-FCRL6 polyclonal Abs. (**C**) Analysis of FCRL6 expression by bone marrow (BM) cells from adult BALB/c mice co-stained with the indicated myeloid, lymphoid, and B lineage differentiation markers and anti-FCRL6 (1C3). Number adjacent to the gate indicates the percentage of CD19^+^FCRL6^+^ lymphocytes. Macrophage (Mϕ), dendritic cells (DC), and natural killer (NK) cells.

**Figure supplement 2.**
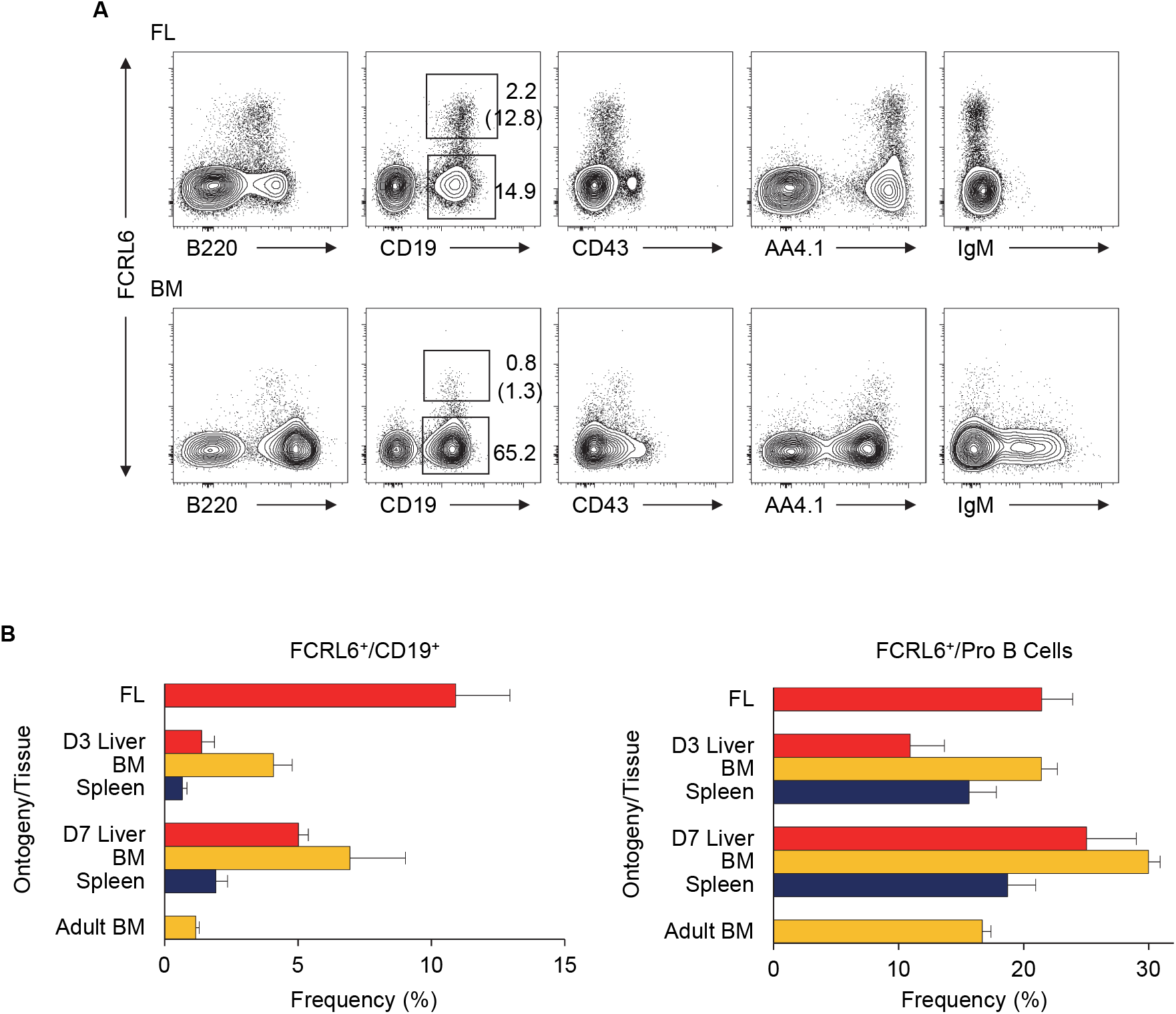
FCRL6 is expressed by B cell progenitors throughout ontogeny. (**A**) Analysis of FCRL6 expression by E18 fetal liver (FL) and adult BM from BALB/c mice co-stained with B lineage differentiation markers and anti-FCRL6 (1C3). Frequencies are indicated adjacent to the gates. The numbers in parentheses indicate the percentage of FCRL6^+^/CD19^+^ lymphocytes. (**B**) Frequencies of FCRL6^+^ cells among total B (gated as in **A** above) and pro B cells (gated as in Figure 1B) determined by staining E18 FL (*n* = 6), adult BM (*n* = 6), and indicated tissues from 3 (*n* = 7) or 7 (*n* = 4) day old neonates. Small horizontal lines indicate the mean (± s.e.m.).

**Figure supplement 3.**
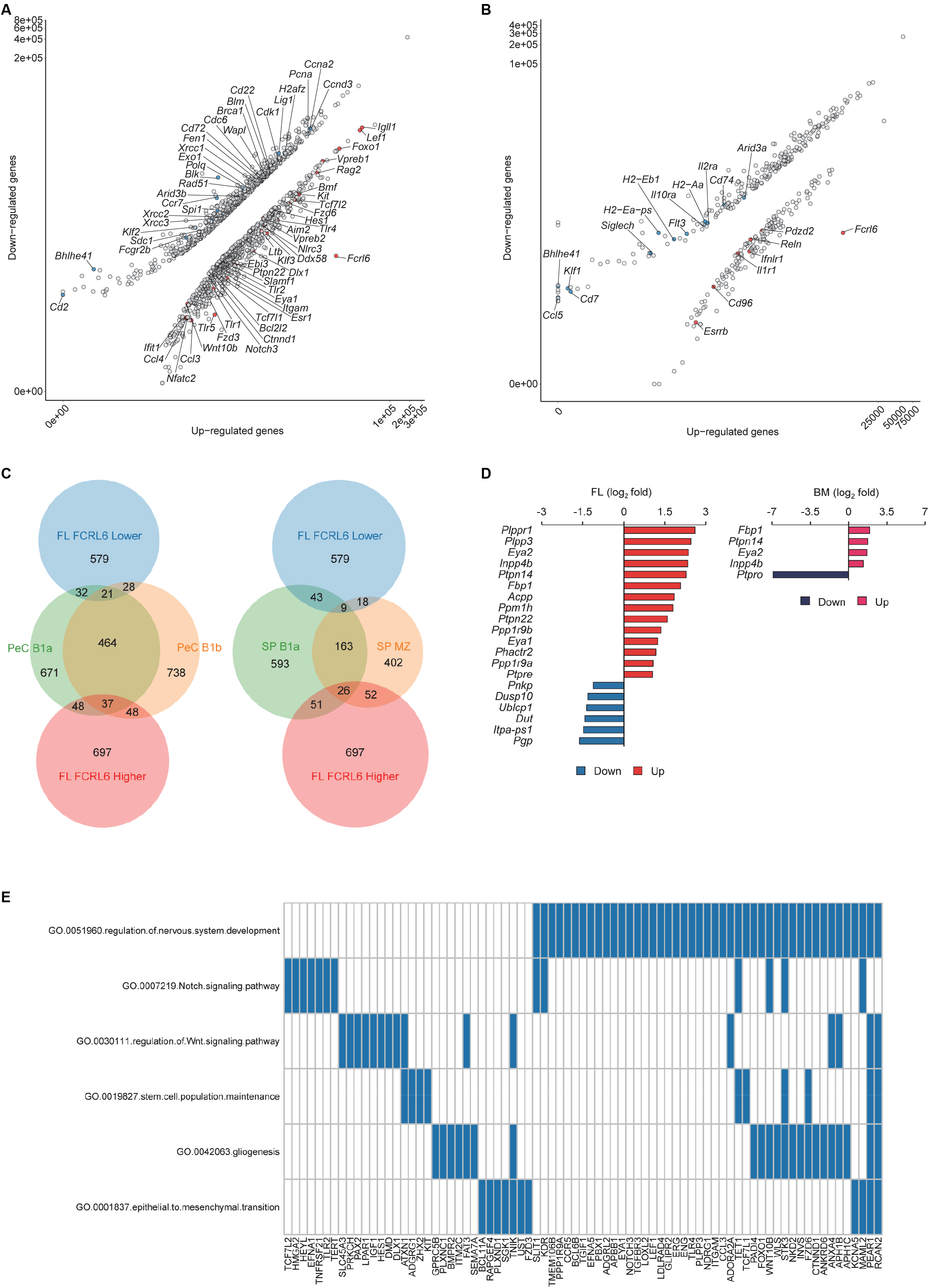
Biologic discrimination of FCRL6^+^ FL and BM pro B cell gene expression profiles. (**A**-**B**) Scatter plots showing differential expression of genes by FL (**A**) and BM (**B**) pro B cells according to up or downregulation of FCRL6 expression. (**C**) Scaled Venn diagrams (https://www.stefanjol.nl/venny) showing gene overlap between up or downregulated DEGs from FCRL6^+^ and FCRL6^-^ FL pro B cells (from Figure 3B) and peritoneal cavity (PeC) B-1a and B-1b or spleen (SP) B-1a and marginal zone (MZ) B cell subsets extracted from the Immgen database (https://www.immgen.org). Upregulated genes (>1.4 fold) from the indicated innate-like B cell subsets were determined relative to follicular B cells from respective tissues using the Immgen population comparison database. (**D**) DEGs encoding phosphatases by FCRL6^+^ FL and BM pro B cells. (**E**) Correlation matrix detailing FL DEGs upregulated by the regulation of nervous system development GO pathway (GO:0051960) that are shared with related pathways.

**Figure supplement 4.**
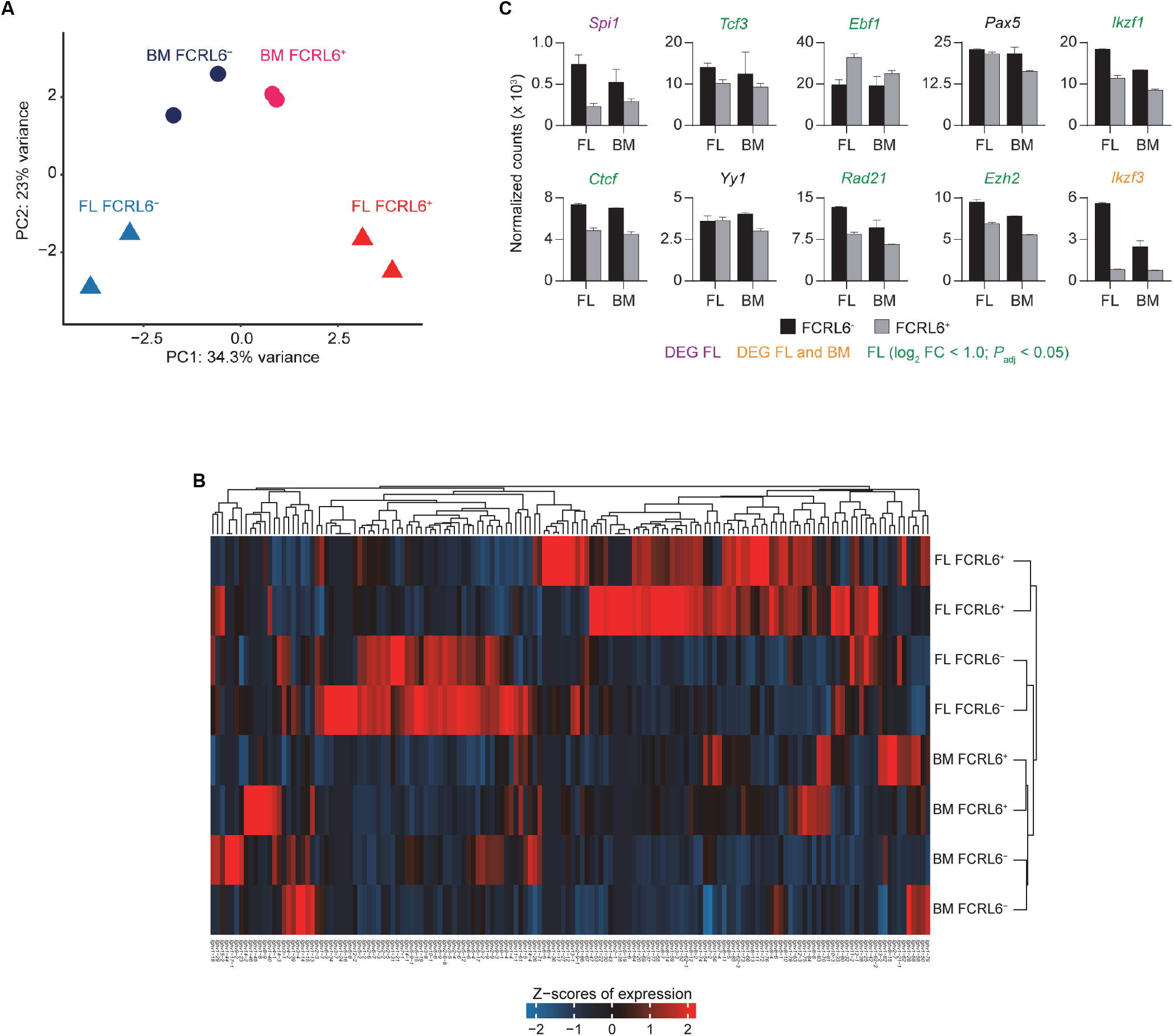
Analysis of *Ighv* repertoire and transcription/accessibility factor gene expression by pro B cell subsets. (**A**) PCA plot of *Ighv* segments expressed by the four pro B cell subsets from FL and BM according to FCRL6 status determined by comparing normalized counts by RNA-seq analysis in duplicate. (**B**) Heat map of *Ighv* segments determined by unsupervised Euclidian clustering. (**C**) Normalized transcript expression of selected transcription and regulatory factors that modulate locus accessibility. Small horizontal lines indicate the mean (± s.e.m.). Note genes labeled in green are significant in FL (*P*_adj_ < 0.05), but did not reach the DEG threshold (one-fold change in log_2_ value).

**Figure supplement 5.**
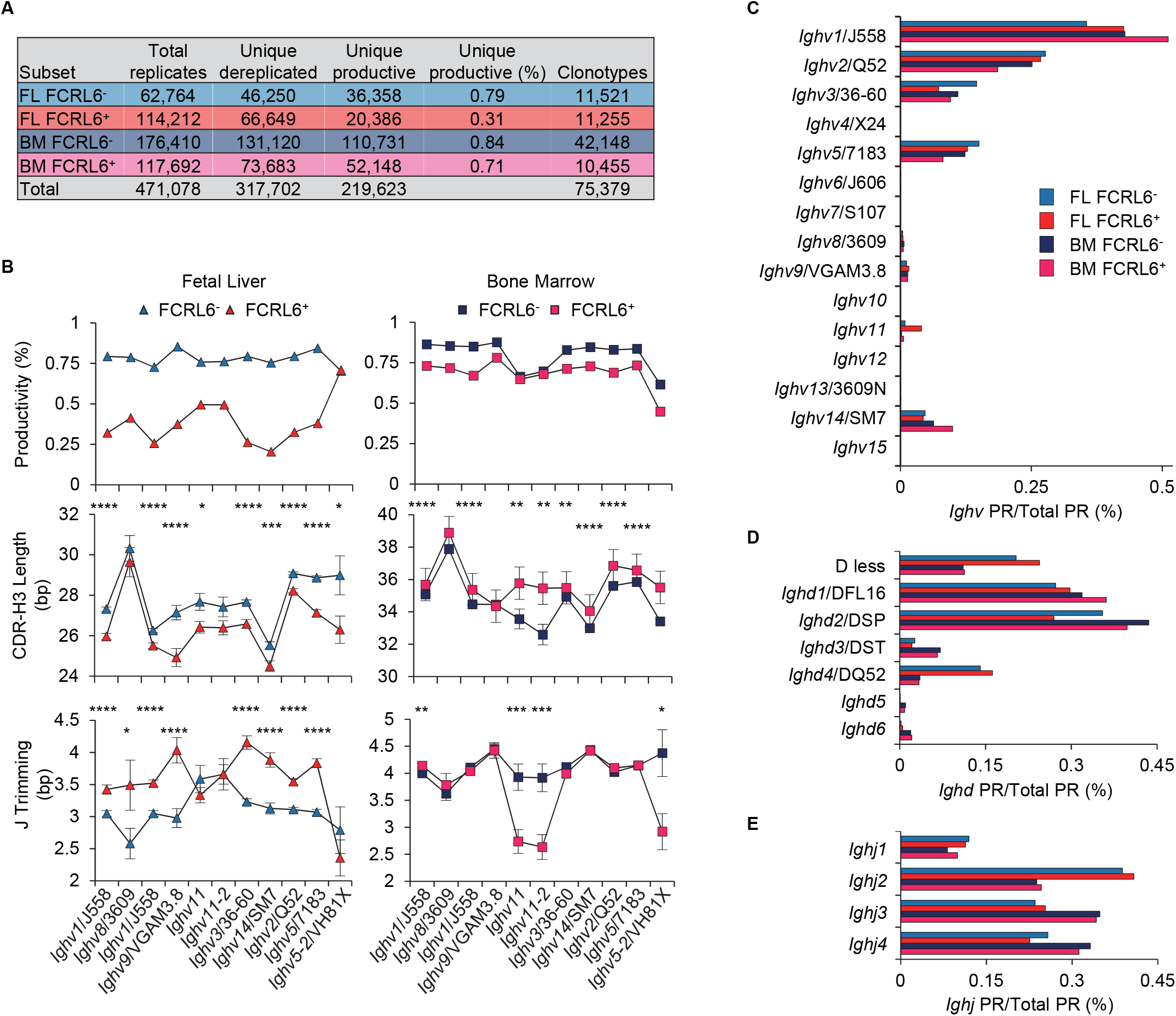
*Ighv* repertoire analysis of FL and BM pro B cells according to FCRL6 status. (**A**) Summary of sequence numbers and characteristics from the four subsets quantitated by replication status and productivity. Clonotypes were defined by identical VJ and CDR-H3 length and ≥90% bp identity. No *Ighv7* sequences were amplified and for *Ighv4*, *6*, *10*, *12*, *13*, and *15,* <100 total dereplicated sequences were identified. (**B**) Productivity, CDR-H3, and J trimming features of unique dereplicated productive sequences of eight *Ighv* families and notable segments (*11-2* and *5-2*) according to locus position. The *Ighv1*/J558 family in (**B**) was segregated according the location of segments within the *Ighv* locus that derive from domains 3 or 4 as in Figure 4A. Small horizontal lines indicate the mean (± s.e.m.). \**P* < 0.05, \*\**P* < 0.01, \*\*\**P* < 0.001 and \*\*\*\**P* < 0.0001 as determined by paired Student’s *t*-test. (**C**-**E**) Frequencies of productive *Ighv* (**C**), *Ighd* (**D**), and *Ighj* (**E**) genes relative to total productive reads (PR) among the four pro B cell subsets.

**Figure supplement 6.**
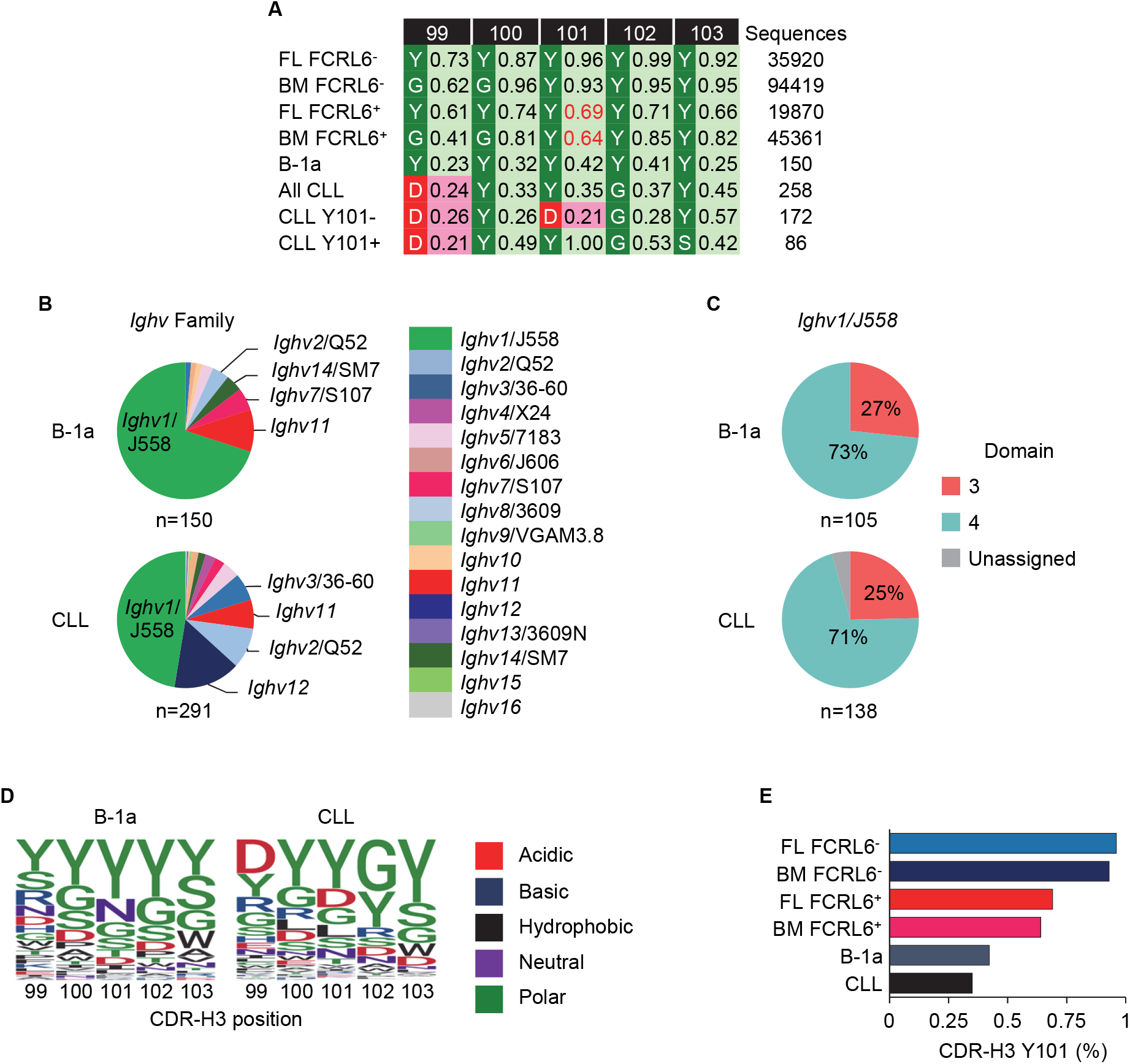
Analysis of *Ighv* usage and CDR-H3 composition from FCRL6^+^ and FCRL6^-^ FL and BM pro B cells, B-1a, and CLL *Igh* sequences. (**A**) Summary of amino acid usage frequency by CDR-H3 position for the four pro B cell subsets, B-1a cells (Yang et al., 2015 – **Table supplement 1**), and CLL expansions (total, Y101 or non-Y101) from published sequences and unpublished CLL sequences (**Table supplement 2**). The total numbers of encoded CDR-H3 sequences analyzed for each subpopulation are indicated in the right column. (**B**) *Ighv* usage for B-1a and CLL sequences by family. The numbers of sequences analyzed are shown below the pie charts. (**C**) *Ighv1*/J558 family usage from B-1a and CLL sequences segregated by *Ighv* locus domain. (**D**) Logo plots detailing the probability of amino acid usage by CDR-H3 amino acid position for B-1a and CLL sequences. (**E**) Relative frequency of Y101 among productive unique sequences (enumerated in **A**) from the four FCRL6^+^ and FCRL6^-^ pro B cell subsets, as well as B-1a cells, and CLL expansions.

**Figure supplement 7.**
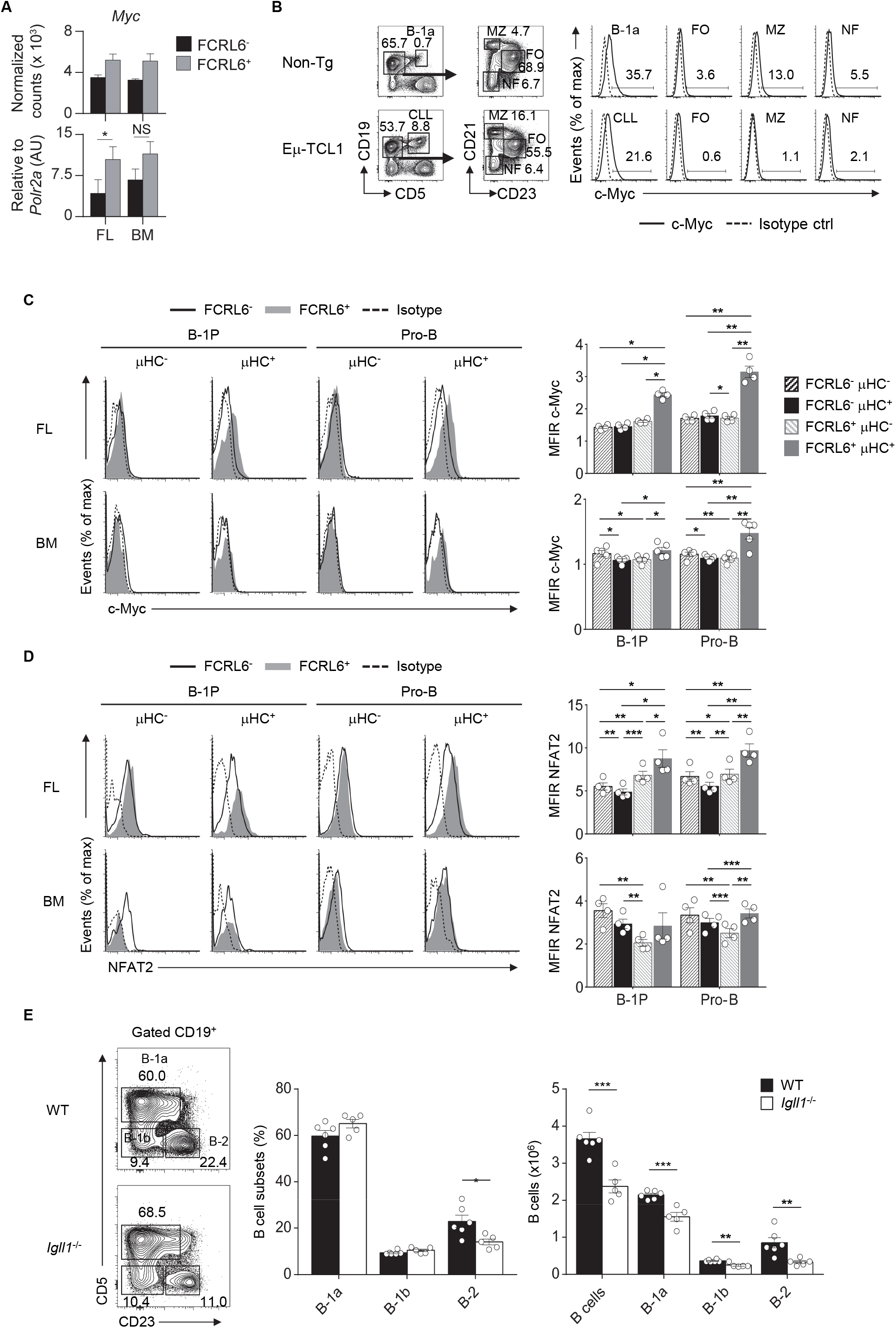
Progenitor B cell transcription factor expression and PeC B cell constitution in λ5 KO mice. (**A**) RNA-seq (above) and RQ-PCR (below) analysis of *Myc* expression by FCRL6^+^ and FCRL6^-^ pro B cell subsets. *Myc* transcripts by RNA-seq (FL 0.57 x log_2_; *P*_adj_ = 0.01) represent normalized mean values from samples in duplicate and by RQ-PCR from three individually sorted samples performed in duplicate. (**B**) Intracellular staining of c-Myc in splenocyte subsets from C57BL/6 and Eμ-TCL1 Tg mice and analysis by flow cytometry. Numbers indicate the percentage of gated cells. Intracellular co-expression of c-Myc (**C**) or NFAT2 (**D**) and μHC in progenitor B cell subsets from FL and BM. (**E**) Analysis of PeC B cell subsets from WT BALB/c and *Igll1*^-/-^ mice. Numbers adjacent to gates indicate frequencies. Each symbol represents an individual mouse. Small horizontal lines indicate the mean (± s.e.m.). \**P* < 0.05; \*\**P* < 0.01 and \*\*\**P* < 0.001 as determined by paired (**A**, **C**-**D**) or unpaired Student’s *t*-test (**E**). Data are representative of at least two independent experiments (**B**-**E**).

**Figure supplement 8.**
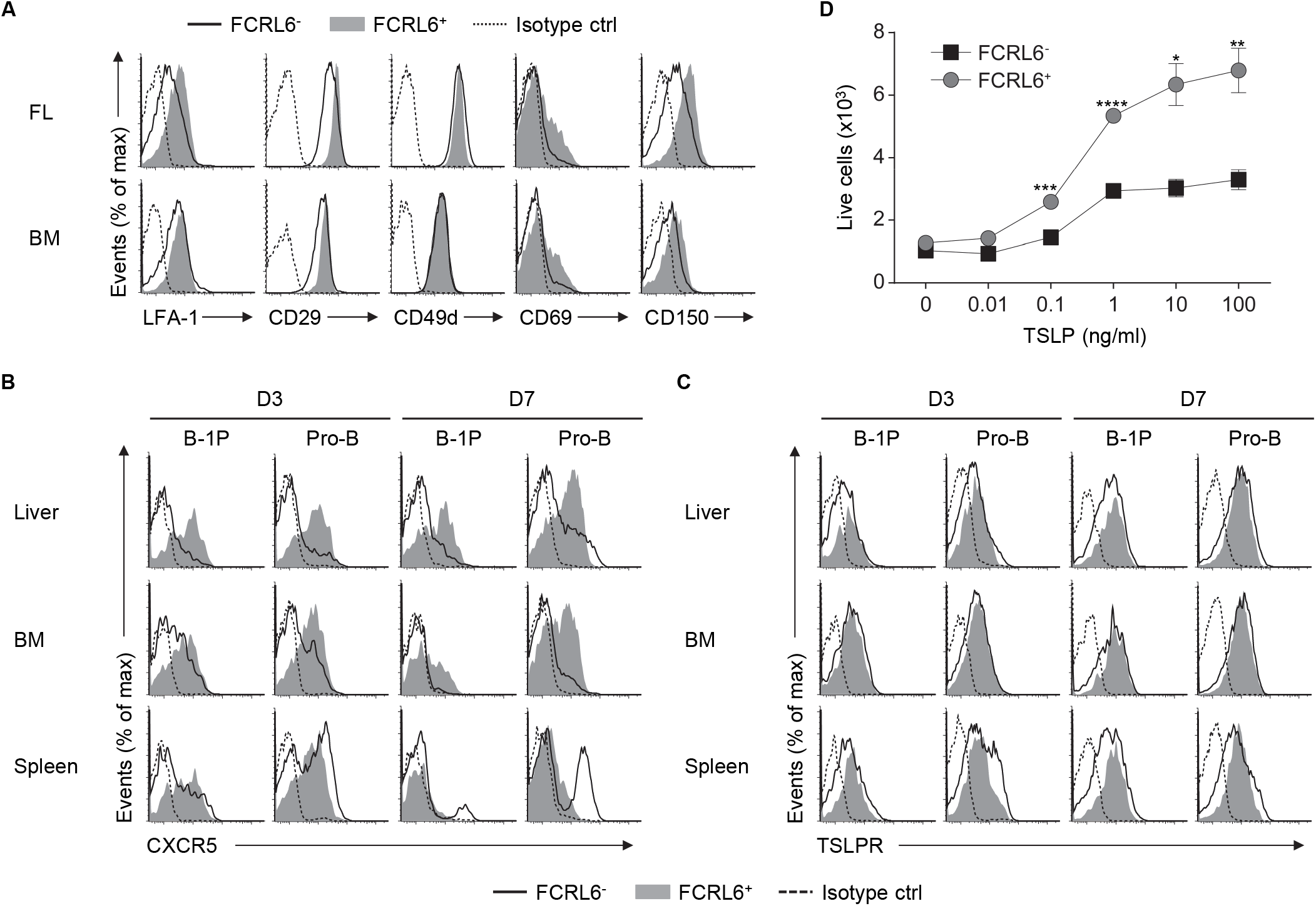
Expression of migration, adhesion, and differentiation factors by FCRL6^+^ and FCRL6^-^ B cell progenitors. (**A**) Flow cytometry analysis of indicated surface markers from E18 FL and adult BM FCRL6^+^ and FCRL6^-^ BM pro B cells. Flow cytometry analysis of CXCR5 (**B**) and TSLPR (**C**) surface expression by D3 and D7 FCRL6^+^ and FCRL6^-^ progenitor B cell subpopulations from the indicated tissues. (**D**) Dose-dependent survival of sorted FCRL6^+^ and FCRL6^-^ BM pro B cells cultured with or without varying concentrations of TSLP (0-100 ng/ml). Small horizontal lines indicate the mean (± s.e.m.). \**P* < 0.05, \*\**P* < 0.01, \*\*\**P* < 0.001 and \*\*\*\**P* < 0.0001 as determined by paired Student’s *t*-test. Data are representative of at least two independent experiments (**A**-**B**) and three independent experiments (**D**).

